# Longitudinal analysis of antibody titers after primary and booster mRNA COVID-19 vaccination can identify individuals at risk for breakthrough infection

**DOI:** 10.1101/2025.03.01.639360

**Authors:** Hyeongki Park, Naotoshi Nakamura, Sho Miyamoto, Yoshitaka Sato, Kwang Su Kim, Kosaku Kitagawa, Yurie Kobashi, Yuta Tani, Yuzo Shimazu, Tianchen Zhao, Yoshitaka Nishikawa, Fumiya Omata, Moe Kawashima, Toshiki Abe, Yoshika Saito, Saori Nonaka, Morihito Takita, Chika Yamamoto, Hiroshi Morioka, Katsuhiro Kato, Ken Sagou, Tetsuya Yagi, Takeshi Kawamura, Akira Sugiyama, Aya Nakayama, Yudai Kaneko, Risa Yokokawa Shibata, Kazuyuki Aihara, Tatsuhiko Kodama, Akifumi Kamiyama, Tomokazu Tamura, Takasuke Fukuhara, Kenji Shibuya, Tadaki Suzuki, Shingo Iwami, Masaharu Tsubokura

## Abstract

A key issue in the post-COVID-19 era is the ongoing administration of COVID-19 vaccines. Repeated vaccination is essential for preparing against currently circulating and newly emerging SARS-CoV-2 variants while enabling people to continue with daily life. Optimizing vaccination strategies is crucial to efficiently manage medical resources and establish an effective vaccination framework. Therefore, it is important to quantitatively understand vaccine-induced immunity dynamics and to be able to identify poor responders with lower sustained antibody titers as potential priorities for revaccination. We investigated longitudinal antibody titer data in a cohort of 2,526 people in Fukushima, Japan, from April 2021 to November 2022 for whom basic demographic and health information was available. Using mathematical modeling and machine learning, we stratified the time-course patterns of antibody titers after 2 primary doses and 1 booster dose of mRNA COVID-19 vaccines. We identified 3 notable populations, which we refer to as the durable, the vulnerable, and the rapid-decliner populations, approximately half of which remained in the same population after the booster dose. Notably, the rapid-decliner population experienced earlier infections than the others. Furthermore, when comparing IgG(S) titers, IgA(S) titers, and T-spot counts between participants who experienced breakthrough infections after booster vaccination and those who did not, we found that IgA(S) titers were significantly lower in breakthrough infected participants during the early stage after booster vaccination. Our computational approach is adaptable to various types of vaccinations. This flexibility can inform policy decisions on vaccine distribution to enhance immunity both in future pandemics and in the post-COVID-19 era.

## Text

The rapid development and distribution of vaccines have played a significant role in controlling the COVID-19 pandemic [1] as we transition towards the post-pandemic era. However, despite this progress, it is crucial to recognize that the risk of COVID-19 has not been completely eradicated, as evidenced by continuous emergence of new SARS-CoV-2 variants [2, 3]. Therefore, alongside fundamental infection control measures such as avoiding crowded places and improving personal hygiene through ventilation and other means [4], vaccination remains pivotal in controlling COVID-19 [5, 6]. In contrast, vaccinating the entire population is unrealistic, leading to the need for a prioritization strategy for additional doses. Notably, the World Health Organization (WHO) recommendations since March 2023 have shifted away from universal COVID-19 vaccine recommendations for moderate- to low-risk populations [7]. This highlights the necessity for new vaccination programs suitable for the post-COVID-19 era that are distinct from the vaccine rollout programs designed to strongly control the pandemic. Addressing this issue requires a quantitative understanding of individual-level immune dynamics after (booster) vaccination, which are distinct from that at the population level. It is critical to identify vulnerable populations and establish priority candidates within these groups [8].

To contribute to this endeavor, we used and evaluated the longitudinal antibody measurements obtained from over 2500 participants who received the Pfizer BNT162b2 or Moderna mRNA-1273 vaccine within the Fukushima vaccination cohort from April 2021 through November 2022 [9]. Our analysis revealed that vaccine-induced antibody dynamics following the primary 2 doses (i.e., primary vaccination) are highly heterogeneous, demonstrating substantial individual variation [10]. Previous research has also reported that additional doses (i.e., booster vaccinations) can rapidly induce high antibody titers [11], even in groups with initially low induced antibody levels (e.g., elderly individuals [12], immunocompromised individuals [13], and those with underlying health conditions [14, 15]). However, it remains unclear how the patterns of vaccine-elicited antibody dynamics after additional doses differ across groups (i.e., individual variation in vaccine-elicited antibody dynamics after additional doses). It is also unclear how these patterns differ from those observed after the primary doses, particularly during the decay phase of the antibody titers. Notably, it is crucial to understand whether individuals classified as “poor responders” who exhibit low sustained antibody titers (i.e., vulnerable and rapid-decliner populations) remain vulnerable or transition to “good responders” who display high sustained antibody titers (i.e., durable population) (and vice versa) after receiving additional doses. Furthermore, individual variations in humoral immune dynamics, their relationship with cellular immune responses, and their connection to the risk of breakthrough infection remain poorly understood. This knowledge is essential for developing effective post-COVID-19 vaccination programs. To gain these valuable insights, we performed a comprehensive analysis by integrating data accumulated within the ongoing Fukushima vaccination cohort, spanning from April 2021 through November 2022.

## Results

### The Fukushima vaccination cohort

Our vaccination study, known as the Fukushima vaccination cohort, was initiated in April 2021 and involved individuals from predominantly rural areas: Soma City, Minami Soma City, and Hirata village in Fukushima, Japan [9, 16–20]. Data for this research were collected from April 2021 to November 2022. The participants included general residents, residents of long-term care facilities, health care workers, frontline workers, and administrative officers. A total of 2526 participants who had received either the Pfizer BNT162b2 or Moderna mRNA-1273 vaccine were recruited (see **Fig 1AB, Supplementary Fig 1**, and **Methods** for further details). The basic demographic and health information of the Fukushima vaccination cohort stratified by gender are provided in **Table 1**.

**Figure 1.**
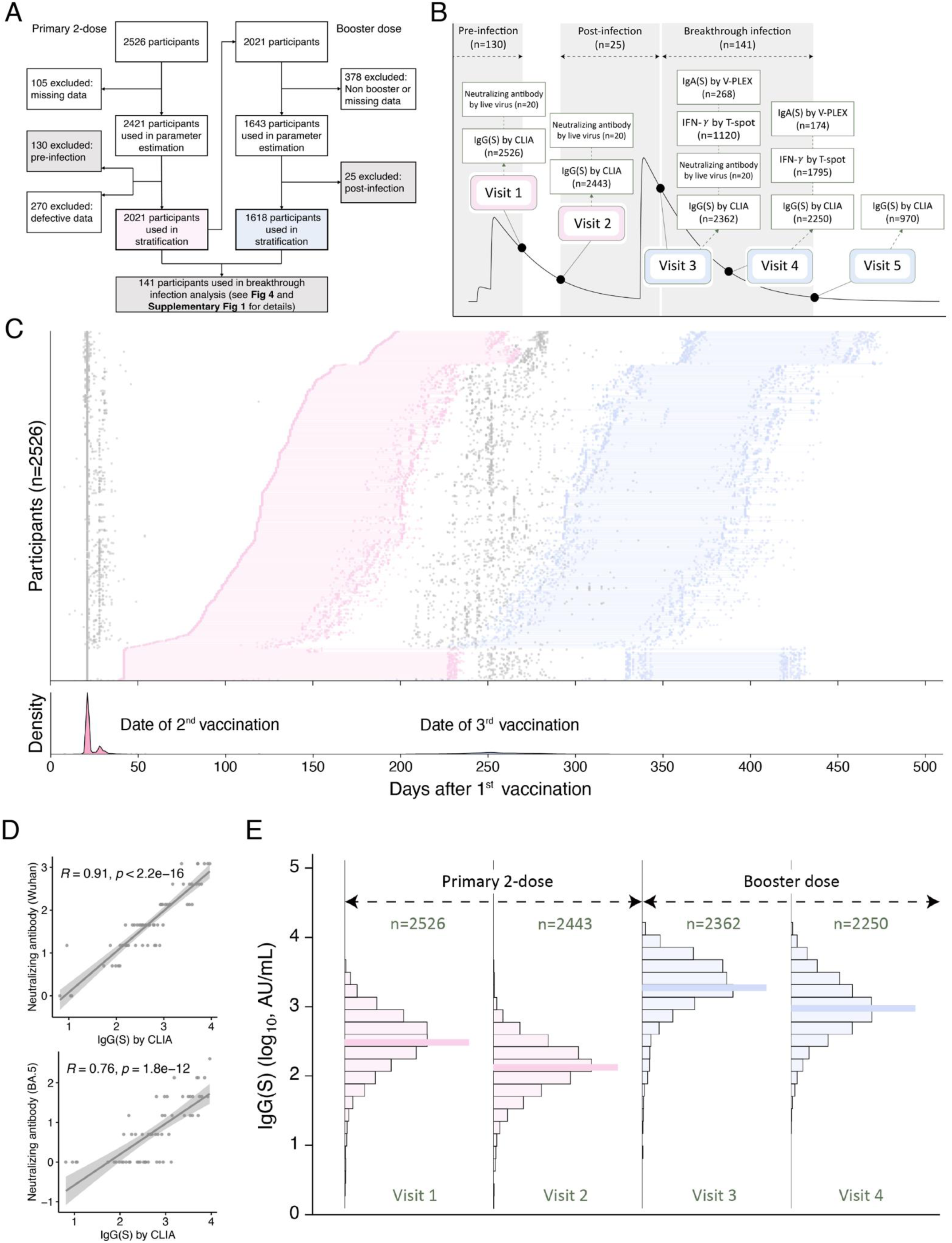
Multi-modal datasets in the Fukushima vaccination cohort: **(A)** The flowchart illustrates the vaccination cohort, including the number of participants, testing procedures, and inclusion criteria for our analysis. Participants with a history of infection (termed ‘pre-infection’ or ‘post-infection’) were analyzed separately (for more details, see **Supplementary Fig 1**). **(B)** The schedule and number of multi-modal datasets obtained through various measurement types at each visit (i.e., blood sampling) are presented. **(C)** A timeline of sample collection from visit 1 to visit 4, primarily used to characterize individual vaccine-induced immunity for each cohort participant, is outlined. The study included 2,526 participants, with a total of 35,277 samples collected. The gray circles indicate the timing of vaccinations, while the pink and blue distributions in the bottom panel represent the dates for the 2nd dose of the primary vaccine and booster vaccinations, respectively. The blue and pink circles indicate blood samplings before and after the booster vaccination, respectively. **(D)** The correlations between IgG(S) and neutralization antibody titers for Wuhan and BA.5 strains by live virus experiments, using the same samples, are depicted. Each data point represents an individual sample, and the correlations were calculated as the Pearson’s correlation coefficients. **(E)** The distribution of longitudinal IgG(S) measurements, obtained through CLIA, is separately plotted based on the timing of blood sampling.

**Table 1.**
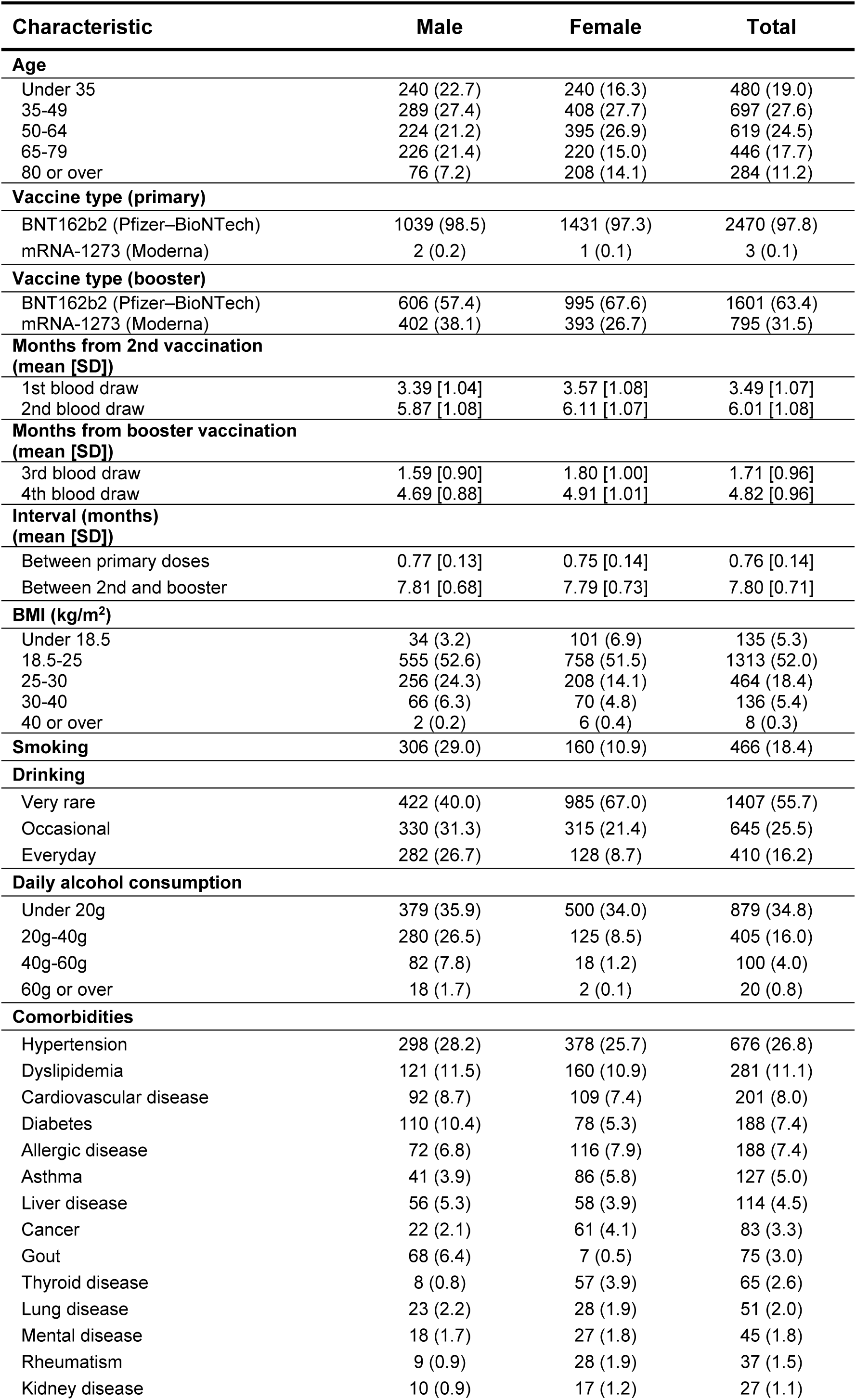
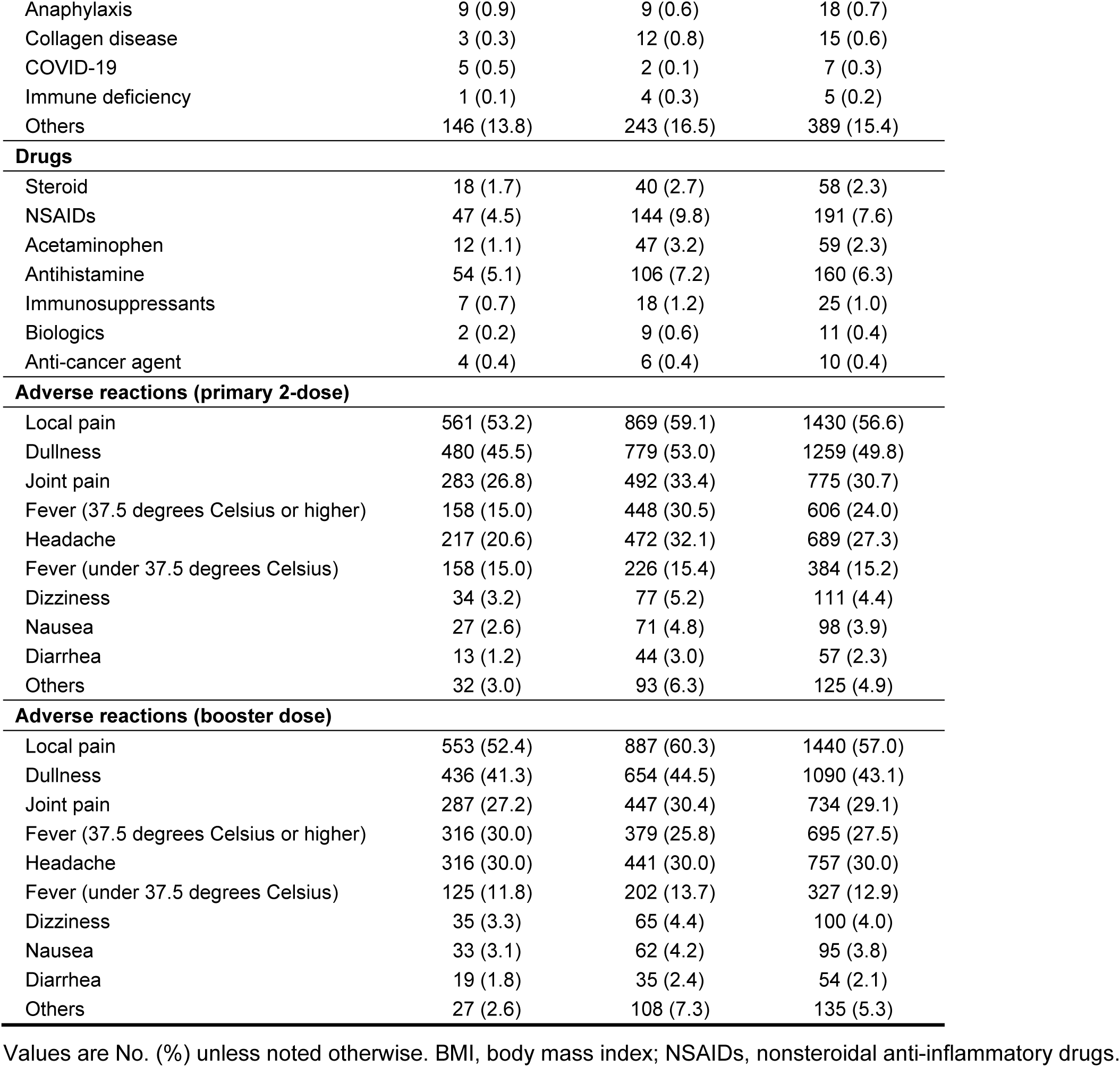
Basic demographics for the Fukushima vaccination cohort.

We conducted a longitudinal analysis of antibody titers in the Fukushima vaccination cohort. For most participants, serum samples were primarily collected at 4 time points (visits 1 through 4), as shown in **Fig 1B**. These antibody titer measurements at the 4 visits were primarily used to characterize individual vaccine-induced antibody dynamics through a mathematical model, which will be discussed later. For some participants, serum samples were also collected prior to visit 1, resulting in a total of 5 time points (see **Methods** for further details). The collection period spanned approximately 3 to 8 months after the 2nd primary dose and around 1 to 10 months after the booster dose of mRNA vaccine. The detailed individual timeline of the sample collections is described in **Fig 1C**.

To quantitatively assess vaccine-elicited immune responses in a high-throughput manner, we conducted a total of 11,759 IgG(S), 11,759 neutralization activity, and 11,759 IgG(N) assays by chemiluminescent immunoassay (CLIA) [21, 22] (as shown in **Fig 1B** and described in the **Methods**). We also conducted live virus experiments using Wuhan and Omicron BA.5 strains to measure neutralizing antibody titers (**Fig 1B** and **Methods**). We compared these titers with IgG(S) measured through CLIA, because previous studies have demonstrated a correlation between neutralizing antibody titers and vaccine-mediated protection (i.e., vaccine efficacy), even against variants of concern [23–26]. Our results indicate a high correlation between these 2 measurements, with correlation coefficients of 0.91 and 0.76 (p-values less than 1.8 × 10^−12^ for both cases) for Wuhan and BA.5, respectively (**Fig 1D**). Consequently, for the evaluation of COVID-19 vaccine-elicited antibody responses, we subsequently utilized longitudinal data of IgG(S) titers against the ancestral strain (i.e., Wuhan strain) from the same individuals as a biomarker for vaccine-elicited humoral immunity (**Fig 1E**). Additionally, for some participants at visits 3 and 4, we assessed cellular immunity after booster vaccination using the T-SPOT.COVID test (Oxford Immunotec). We also measured IgA(S) levels in participants with breakthrough infections and in control subjects to evaluate the risk of breakthrough infection following booster vaccination at visits 3 and 4 (see **Fig 1B**, **Supplementary Fig 1**, and **Methods**).

Our aim was to investigate and characterize the variations in the immune dynamics to primary and booster COVID-19 vaccination at the individual level. Furthermore, we quantitatively examined these responses, focusing particularly on individuals who were poor responders. The following sections provide valuable insights to prepare an effective vaccination strategy for the post-COVID-19 era.

### Stratifying primary and booster vaccine-elicited antibody dynamics

As illustrated by the distribution of antibody titers induced by the primary vaccinations in **Fig 1E**, the average IgG(S) titer declined from 457.5 to 192.4 arbitrary units (AU)/mL — equivalent to a 2.4-fold decrease — over approximately 3 months at the population level (i.e., from visit 1 to visit 2). This finding aligns with previous studies [27–30]. In contrast, after the booster vaccination, titers increased significantly. Subsequently, the titers declined from 2739.4 to 1496.2 AU/mL on average (i.e., 1.8-fold decrease) 5 months after the booster vaccination (i.e., from visit 3 to visit 4). While these results demonstrate that a booster vaccination leads to a significant increase in the antibody response, followed by a subsequent decline at the population level, the specific quantitative dynamics of antibodies at the individual level, particularly in terms of individual variations, remain unclear.

To better understand this individual heterogeneity, we applied our recently developed mathematical model describing vaccine-elicited antibody dynamics [10, 31]. We fully reconstructed the dynamics of IgG(S) titers at the individual level over a period of approximately 500 days after the 1st vaccination (visits 1 through 4) for all individuals in the Fukushima vaccination cohort in **Supplementary Fig 2A** (see **Methods** in detail). To validate our mathematical model and the reconstructed antibody titer curves, we further collected independent blood samples (which were not used in the fitting) from 113 and 536 individuals after the primary and booster vaccinations, respectively (see **Methods** and **Supplementary Fig 1** for details). Of note, as shown in **Supplementary Fig 2BC**, we confirmed that our reconstructed curves well match the IgG(S) titers measured from these independent samples. Note that individuals with a history of natural infection will be evaluated in the next section (see **Supplementary Fig 8**).

First, we performed an unsupervised clustering analysis using a random forest dissimilarity [32, 33] to stratify the time-course patterns of antibody dynamics elicited by the primary vaccination into 6 groups (i.e., G1 to G6) (see **Methods** and **Supplementary Fig 3A-D** in detail). **Fig 2A** represents a two-dimensional Uniform Manifold Approximation and Projection (UMAP) embedding of these 6 groups, which clearly shows that G5 and G6 are separated from the other groups. Using a different color for each group, we also plotted the reconstructed individual antibody dynamics as shown in the left panel of **Fig 2B** (see also **Supplementary Fig 3D**). The projected time course of each group showed that sustained antibody titers remained high among individuals in G1 (N=163), whereas those in G6 (N=296) were low (the right panel of **Fig 2B** and **Supplementary Fig 3E**). We classified G1 as a “durable” population and G6 as a “vulnerable” population in our analysis. In contrast, individuals in G4 (N=272) exhibited an atypical pattern (called “rapid-decliner”), characterized by a high peak of antibody titers that rapidly decreased to a very low level (**Fig 2B** and **Supplementary Fig 3DE**). We considered G1 as “good” responders, and G6 and G4 as “poor” responders. The other groups, G2 (N=157), G3 (N=849) and G5 (N=284), showed intermediate dynamics and were considered as intermediate responders.

**Figure 2.**
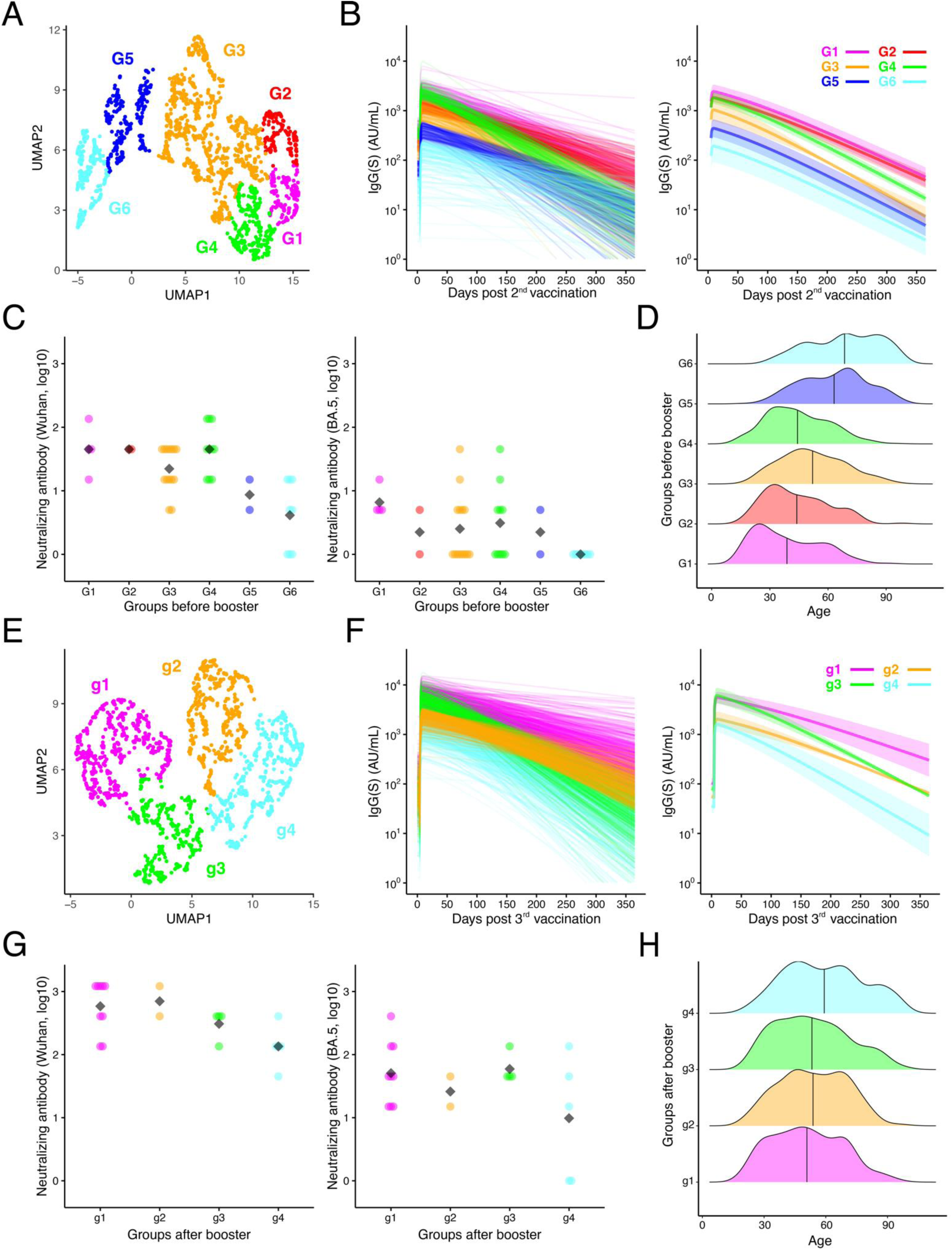
Primary and booster vaccine-elicited humoral immune responses: **(A)** UMAP of stratified antibody response to the primary vaccination, based on the extracted features from the reconstructed individual-level antibody dynamics and estimated parameters (see **Methods** for details), is shown. Data points represent each participant and are colored according to groups (G1 to G6). **(B)** The reconstructed individual antibody dynamics in each group G1 to G6 (left), and their time-course patterns highlighted by the Partial Least-Squares Discriminant Analysis (PLS-DA) (right), are shown. The solid line and shaded area in the figure on the right indicate mean and mean ± standard deviation of rearranged IgG(S) values by PLS-DA for each time, respectively. **(C)** The neutralizing antibody responses for the Wuhan (left) and BA.5 (right) strains, quantified by live virus experiments using a subset of blood samples measured at visit 1 (left) and visit 2 (right) belonging in each group, are shown, respectively. **(D)** The age distributions of participants belonging in each group are shown. **(E)**, **(F)**, **(G)**, and **(H)** show the results of the same analysis described in **(A)**, **(B)**, **(C)**, and **(D)** using post-booster stratified groups (i.e., g1 to g4) instead of G1 to G6. Note that **(G)** is based on a subset of blood samples collected at visit 3 (left) and visit 4 (right).

Additionally, we compared the neutralizing antibody titers for the Wuhan and BA.5 strains through live virus experiments, measured at visit 1 and visit 2, across the stratified groups (**Fig 2C**). Consistent with the IgG(S) titer dynamics, the neutralizing titers against the Wuhan strain were highest in the durable (G1) population and lowest in the vulnerable (G6) population. In contrast, only durable individuals exhibited relatively high neutralizing titers against the Omicron strain (higher than those against the Wuhan strain among vulnerable individuals), while the other groups displayed very low neutralizing titers, particularly among the vulnerable populations (almost 0). These results further support the durability and vulnerability of G1 and G6 for the “unknown” variants of concern because the Omicron subvariant BA.5 had not emerged at the time of blood sampling in the Fukushima vaccination cohort (the main blood sampling period was from September 2021 to June 2022) [34, 35]. We also examined the age dependency of the 6 groups (**Fig 2D**): G1, G2, G3, G4, and G5 primarily consisted of individuals aged 35 to 60 years (young and middle-aged) and G6 consisted of individuals aged 52 to 80 years (middle-aged and elderly), respectively. It is worth noting that the vulnerable population (G6) includes individuals who may not necessarily be older adults, and the rapid-decliners (G4) primarily comprised participants aged 32 to 56 years at the time of primary vaccination (discussed later).

In a similar manner, we stratified the time-course patterns of antibody dynamics following the booster vaccination into 4 groups (referred to as g1 to g4, called post-booster stratified group) (see **Fig 2E** and **Methods** for details). We confirmed that individuals in g1 (N=478) represented the durable population, while those in g4 (N=446) were classified as vulnerable. Additionally, individuals in g3 (N=313) were identified as rapid decliners (**Fig 2F** and **Supplementary Fig 3FG**). We considered g1 as “good” responders, g4 and g3 as “poor” responders, and g2 (N=381) as intermediate responders. When comparing the peaks of antibody titers induced by the primary and booster vaccinations (**Supplementary Fig 3EG**), it is evident that the booster vaccination robustly elicits antibody titers in all individuals, regardless of their initial antibody titers or the stratified groups G1 to G6 (e.g., [36–38]). The fold increase in antibody titers due to the booster vaccination, as calculated in **Supplementary Fig 4**, further confirms this significant induction.

We also compared the booster vaccine-elicited neutralizing antibody titers against both the Wuhan and BA.5 strains, measured at visit 3 and visit 4, among the stratified groups g1 to g4 (**Fig 2G**). The durable population (g1) exhibited high neutralizing antibody titers against both strains, while even the vulnerable population (g4) regained neutralizing capability against the Omicron strain BA.5. Interestingly, we observed only small differences in the age distribution among the stratified groups after the booster vaccination (**Fig 2H**): g1, g2, g3, and g4 primarily consisted of individuals aged 37 to 64 years, 42 to 67 years, 38 to 66 years, and 44 to 73 years, respectively. Compared with the case for the primary vaccination (**Fig 2D**), age dependency appears to be weak for the antibody titers elicited by the booster vaccination.

To compare the longevity of primary and booster vaccine-induced antibody titers, we analyzed the proportions of individuals with IgG(S) antibody titers below predefined thresholds (i.e., 10 and 200 AU/mL: see **Supplementary Fig 5** in detail) in each stratified group (G1 to G6 and g1 to g4) at 180 and 365 days after the primary and booster vaccinations. Our simulations also revealed that the majority of G1/G2 and g1/g2 maintained antibody levels above 200 AU/mL for at least 6 months. Moreover, approximately 65% of g1 still displayed no decline below 200 AU/mL after 1 year following the booster vaccinations (see **Supplementary Fig 5**). In contrast, the vulnerable populations (i.e., G6 and g4) exhibited a low induction of antibody titers after vaccination and experienced a rapid decline in efficacy (i.e., less than 10 AU/mL). Roughly 57% and 40% of individuals within these populations had very low antibody titers after 1 year, respectively, comparable to those observed in unvaccinated individuals [39]. Individuals in the rapid-decliners group (i.e., G4 and g3) initially demonstrated a high induction of antibody titers following the vaccinations but experienced a rapid decay; approximately 56% and 14% of them showed a reduction in antibody levels below 10 AU/mL at 1 year after the primary and booster vaccinations.

Interestingly, when calculating the proportions of transitions between stratified groups resulting from the primary and booster vaccinations (i.e., from G1-G6 to g1-g4) in **Supplementary Fig 6A**, approximately 60% of the durable population observed during the primary vaccination phase statistically significantly shifted to the durable group, g1 (*p* = 3.1 × 10^−13^ by the Pearson’s chi-square test with the Bonferroni correction), while the other 40% shifted to nondurable groups (i.e., g2-g4). Additionally, 55% and 36% of the vulnerable and rapid-decliner populations statistically significantly remained in the same populations even after receiving the booster vaccinations (*p* = 2.2 × 10^−16^ and *p* = 2.4 × 10^−10^, respectively). These group transitions are visualized in **Supplementary Fig 6B** by plotting participants in G1-G6 on a UMAP plane for g1-g4 generated by using the features of antibody dynamics after the booster dose described in **Fig 2E**.

### Influence of infection history on stratified vaccine-elicited antibody dynamics

We placed individuals with a history of natural infections prior to booster vaccination into 2 groups: (1) individuals with a history of infection before visit 1, who were likely infected before receiving the primary 2 doses (“pre-infection”, N=130), and (2) individuals with a history of infection between visit 2 and visit 3, who were likely infected after the primary 2 doses but before the booster dose (“post-infection”, N=25) (see **Fig 1AB** and **Supplementary Fig 1AB**). Although we lack sequence information to definitively determine the specific variants involved in our study, given the timing of infection for the recovered cases and the prevalent strain in the Fukushima vaccination cohort, we expect that the variants observed were either Wuhan or Alpha variants.

To understand how these infection histories contributed to the stratification of primary and booster vaccine-elicited antibody dynamics, we reconstructed the elicited-antibody dynamics of the pre-infection and post-infection individuals and compared them with the elicited-antibody dynamics of individuals without an infection history. First, when comparing the engineered features (i.e., the peak, duration, and AUC) described in **Supplementary Fig 7A**, we confirmed that the infection history robustly elicits antibody titers in general (**Supplementary Fig 8AC**). Particularly, there are significant differences between individuals with infection histories and those in the vulnerable populations, such as G6 and g4. This is consistent with the priming of immune responses by a history of infection, as discussed in [40–43]. However, we did not observe a significant trend where individuals with infection histories were consistently classified into a specific group, such as the durable populations, G1 and g1 (**Supplementary Fig 8BD**). This indicates that there is significant heterogeneity in the vaccine-elicited antibody titers even among individuals with infection histories. We plan to conduct a more detailed evaluation of individuals with breakthrough infections at a later time.

### Vaccine-elicited cellular immune dynamics within stratified groups

We conducted an extensive analysis of T-spot responses at 2 different time points after administering the booster vaccination (N=1120 and N=1795 at visit 3 and visit 4 of Fukushima vaccination cohort, respectively) (**Fig 1B**). Among them, we used 854 participants (216, 183, 188, and 267 participants in g1, g2, g3, and g4) in the T-spot analysis, who were used in the post-booster stratification analysis and had T-spot data for both of 2 different time points. These samplings were conducted approximately 3 months apart, aiming to assess the vaccine-induced cellular immune responses, specifically focusing on the IFN-γ production (**Fig 1B**) [44]. The paired IgG(S) and T-spot counts were compared, and they were measured from the same blood samples of the same individuals, as described in **Fig 3A**. The results showed weak or no positive correlations, regardless of the timing of sampling. Similar correlations were also confirmed for other features (**Supplementary Fig 9**). These findings align with previous studies [44, 45].

**Figure 3.**
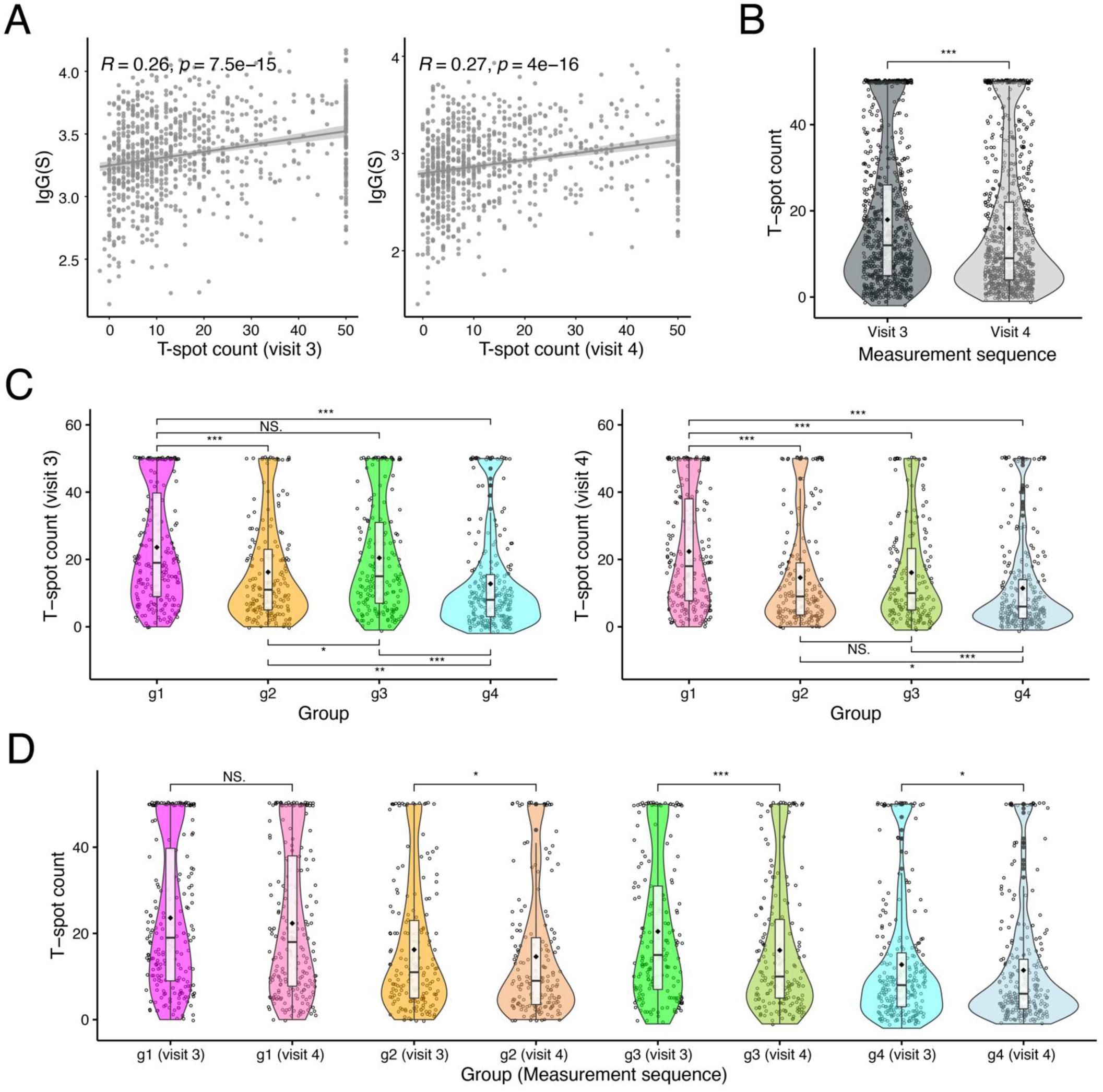
Primary and booster vaccine-elicited cellular immune responses: **(A)** The relation between the T-spot response and IgG(S) measured in the same sample are shown. The left and right plots show the results of linear regressions between the 1st (visit 3) and end (visit 4) measured T-spot counts and the corresponding IgG(S), respectively. The *R* and *p* on the top of each panel indicate the Pearson’s correlation coefficient and its p-value, respectively. **(B)** The distributions of 1st and 2nd measured T-spot counts at visits 3 and 4, respectively, are shown (N=854 for both time points). **(C)** The distributions of 1st (visit 3, left) and 2nd (visit 4, right) measured T-spot counts in each group are shown (N=216, 183, 188, and 267 for g1, g2, g3, and g4). **(D)** The distributions of T-spot counts for all pairs of group and measurement sequence are shown. For **(B)**, **(C)** and **(D)**, statistical significances are calculated by the pairwise Mann-Whitney U test. Also, for **(C)** and **(D),** p-values are corrected by Bonferroni’s method (NS.: p-value > 0.05, ∗: p-value ≤ 0.05, ∗∗: p-value ≤ 0.01, and ∗∗∗: p-value ≤ 0.001, respectively). The statistical significance of the difference in T-spot counts is indicated on the top or bottom of the plots.

As anticipated from the decline in antibody titers as explained in **Fig 2FG** and **Supplementary Fig 3FG**, the average T-spot counts decreased from 17.9 to 15.9 spot-forming cells (SFCs) (i.e., 1.13-fold decrease) over the relatively short period of 3 months at the population level (**Fig 3B**) (i.e., from visit 3 to visit 4). To examine the decay characteristics of T-spot counts in more detail, we plotted and compared T-spot counts within groups g1 to g4, as shown in **Fig 3C**. We observed statistically significant large variations in the strength of booster vaccine-elicited cellular responses, similar to the humoral immune response mentioned above. At approximately 5 months after the booster vaccination at visit 3 (left panel in **Fig 3C**), the T-spot counts among the durable and rapid-decliner populations (i.e., g1 and g3) were the highest (23.6 and 20.5 SFCs on average, respectively), while those among the vulnerable population, g4, were the lowest (12.8 SFCs on average). These findings suggest a potential correlation between the durability and vulnerability of g1 and g4, as defined by the booster vaccine-elicited antibody dynamics, and the cellular immune responses. A similar trend was confirmed after 3 months (i.e., at visit 4), except for the rapid-decliner population, g3 (right panel in **Fig 3C**). While there was no significant difference in the T-spot counts between g1 and g3 at visit 3, a significant difference seemed to emerge at visit 4 (discussed below).

Regarding the longevity of booster vaccine-induced cellular immune responses, we further compared the T-spot counts at different sampling times, at visit 3 and visit 4, within the post-booster stratified groups (**Fig 3D**). Interestingly, unlike the population-level comparisons mentioned earlier (**Fig 3B**), we observed no significant decrease in the T-spot counts among the durable population, g1. This indicates that the booster vaccination effectively primed long-term sustained cellular immune responses and supports the durability of g1. In contrast, the T-spot counts significantly decreased within the non-durable populations (i.e., g2, g3, g4) during the 3 months. Among them, the rapid-decliners showed the largest reduction in T-spot counts (21.3% reduction on average), while the T-spot counts of vulnerable individuals decreased to the lowest level (5.3% reduction on average).

### Relationship between IgG(S) titers and the risk of breakthrough infection

A decrease in antibody titers may increase the risk of breakthrough infection. Especially, the drastic decay characteristics of both IgG(S) titers and T-spot counts induced by vaccination in the vulnerable and rapid-decliner populations (i.e., g4 and g3) indicate that these populations may be at high risk of breakthrough infection. Understanding the relationship between the booster vaccine-elicited antibody dynamics and the risk of breakthrough infection is crucial to design vaccination programs suitable for the post-COVID-19 era.

To investigate the relationship between antibody dynamics and the risk of breakthrough infection after the COVID-19 booster vaccination, we analyzed IgG(S) titers in participants from the Fukushima vaccination cohort who experienced breakthrough infections within approximately 9 months (i.e., infections between the booster vaccination and visit 5) after booster vaccination (**Fig 4A** and **Supplementary Fig 1B**). We identified 141 participants with breakthrough infections after booster vaccination based on measured IgG(N) titers and additional Roche diagnostic tests (see **Fig 4C** and **Methods** for details). All 141 cases were first-time infections (i.e., no prior history of infection) and were either mildly symptomatic or asymptomatic. Of these, 51 participants were infected between visit 3 and visit 4 (first testing positive at visit 4), and 36 of them reported a specific infection date. The remaining 90 participants were infected between visit 4 and visit 5 (first testing positive at visit 5), with 76 reporting a specific infection date. Note that the 90 participants who were infected between visit 4 and visit 5 were included in the stratification analysis after the booster vaccination, whereas the 51 participants infected between visit 3 and visit 4 were excluded, as at least 2 observations (at visit 3 and visit 4) without infection are required to reconstruct antibody dynamics using our mathematical model (**Fig 4AC** and **Supplementary Fig 1AB**).

**Figure 4.**
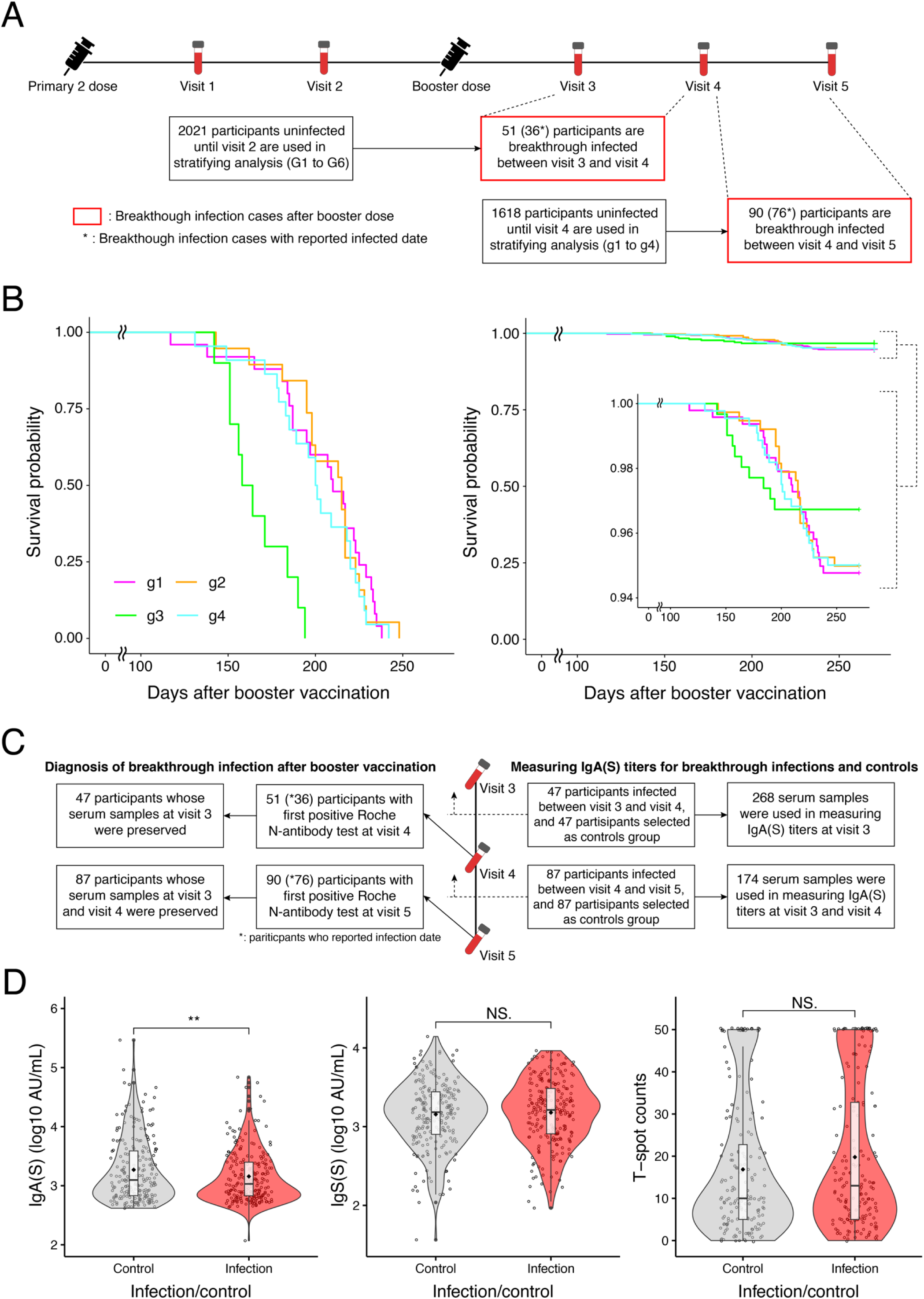
Breakthrough infection after booster vaccination: **(A)** The flowchart illustrates the number of breakthrough infection cases after booster vaccination in the Fukushima vaccination cohort. For more detailed information, please refer to **Supplementary Fig 1AB**. **(B)** The survival probability for each post-booster stratified group, analyzed for breakthrough infection, is shown separately for only the participants with breakthrough infection (left) and for the total population of each group (right). Survival analysis was conducted on the 76 participants who experienced a breakthrough infection after visit 4 and reported a specific infection date. **(C)** The flowchart illustrates the serum samples used for the Roche diagnostic assays (left side) and those used to measure IgA(S) titers (right side), categorized by the timing of infection in breakthrough infection cases. For the control group, IgA(S) titers were measured using the same samples (i.e., samples with the same visit number) as the corresponding breakthrough infection cases. Among the 141 participants (51+90) with confirmed infection by Roche immunoassay, IgA(S) titers were measured in 134 participants (47+87) with available serum samples. **(D)** Comparison of IgA(S) titers, IgG(S) titers, and T-spot counts between the breakthrough infection group and the control group is shown. The statistical analysis was performed using paired t-test (NS.: p-value > 0.05, and ∗∗: p-value ≤ 0.01, respectively).

First, we evaluated the association between the 90 breakthrough infection cases and the post-booster stratified groups (i.e., g1-g4). Among these 90 cases, the distribution across stratified groups g1-g4 was 25, 21, 17, and 27 participants, respectively. Of the 1,618 participants analyzed, 5.6% experienced a breakthrough infection within 9 months of receiving a booster vaccination, with group-specific rates of 5.2%, 5.5%, 5.4%, and 6.1% for groups g1 through g4, respectively. Our findings indicate that the most vulnerable group, g4, had a slightly higher likelihood of breakthrough infection compared with the other groups.

Next, to analyze differences in the timing of breakthrough infections after booster vaccination among the post-booster stratified groups, we conducted a survival analysis for each group (**Fig 4B**). We first analyzed 76 of the 90 breakthrough infection cases with a reported infection date. Two survival analyses were conducted: one focused solely on participants with breakthrough infections (**Fig 4B**, left), and the other on the total population within each stratified group (**Fig 4B**, right). Both analyses used Kaplan-Meier estimation. Notably, participants in the rapid-decliner group (g3) experienced breakthrough infections earlier than those in other groups (p-value < 0.01). In group g3, the probability of survival following a breakthrough infection declined sharply after 150 days after booster vaccination, with all infections occurring within 200 days (**Fig 4B**). This suggests that the timing of breakthrough infections is closely tied to the time-series pattern of IgG(S) titers, making it challenging to predict breakthrough infection risk based on a single IgG(S) titer measurement taken soon after vaccination (discussed further in the next section). Note that we further conducted a survival analysis on a total of 112 participants, which included 36 of the 51 breakthrough-infected participants who reported their infection date, as well as the 76 participants shown in **Fig 4B**. Similar results were observed, as shown in **Supplementary Fig 10**.

### Relationship between serum IgA titers and the risk of breakthrough infection

In addition to IgG(S) response, we analyzed the contributions of other anamnestic immune responses, which are also likely to play a protective role against breakthrough infections. Previous studies have reported that IgA antibodies against the SARS-CoV-2 ancestral strain spike antigen in the nasal mucosa are associated with a reduced risk of breakthrough infection [46]. Also, in our previous study, we reported that nasal secretory anti-spike IgA antibodies reduce viral RNA load and infectivity [47]. To further investigate the relationship between IgA response and the risk of breakthrough infection, we retrospectively measured IgA(S) titers in a subset of the Fukushima vaccination cohort at visit 3 or visit 4 using preserved serum samples.

First, we examined the correlation between nasal and serum IgA antibodies by reanalyzing our original data published in [47]. We observed a strong correlation between the two, with Pearson correlation coefficients of 0.77 for the ancestral strain and 0.75 for the BA.1 strain (**Supplementary Fig 11**). This finding suggests that serum IgA antibodies may serve as an indicator of immune response likely to offer protection against breakthrough infection.

Then, we measured IgA(S) titers using an electrochemiluminescence immunoassay in 134 participants from 141 breakthrough infection cases using serum samples collected at visit 3 or visit 4 (the serum samples were not available for 7 participants), along with an equal number of participants selected as controls (**Fig 4C**). Specifically, using a case-control study approach, the control group was matched to the breakthrough infection cases based on clinical information, stratified group, and timing of data collection after vaccination (see **Methods** for details). IgA(S) titers were measured in serum samples collected after booster vaccination, up to the time of breakthrough infection. For the 134 participants with breakthrough infections, IgA(S) titers were measured from visit 3 samples for 47 participants infected between visit 3 and visit 4, and from both visit 3 and visit 4 samples for 87 participants infected between visit 4 and visit 5. IgA(S) titers for 134 participants in the control group were also measured from serum samples taken at the same visit times as for the corresponding participants in the breakthrough infection group (**Fig 4C**). We measured IgA(S) titers against the ancestral strain and against the BA.1, BA.2, BA.5, BQ.1, and XBB.1 variants and found that the IgA(S) titers against the ancestral strain were highly correlated with those for the other variants, as reported in [46] (**Supplementary Fig 12**). Therefore, we used the ancestral IgA(S) titers for the subsequent analysis without loss of generality.

We analyzed the relationship among IgA(S) titers, IgG(S) titers, and T-spot counts against the ancestral strain using the Pearson correlation coefficient. The correlation coefficients for the relationships between IgG(S) titers and IgA(S) titers and between IgA(S) titers and T-spot counts were 0.52 and 0.11, respectively (**Supplementary Fig 13**). That is, IgA(S) and IgG(S) titers showed a weak positive correlation, whereas a weak or no correlation was observed between IgA(S) titers and T-spot counts. Notably, the correlation between IgA(S) titers and T-spot counts was weaker than that between IgG(S) titers and T-spot counts (**Fig 3A**), suggesting that IgA(S) is more independent of cellular immunity (as measured by T-spot counts) than is IgG(S).

We also compared IgA(S) titers, IgG(S) titers, and T-spot counts between the breakthrough infection group and the control group. Interestingly, we found that IgA(S) titers in the breakthrough infection group were statistically significantly lower than those in the control group, whereas there were no significant differences between the 2 groups for IgG(S) titers and T-spot counts (**Fig 4D**). It is important to note that the control cases were selected from the same stratified group as the breakthrough infection cases, ensuring that participants with similar IgG(S) dynamics were intentionally chosen. As a sensitivity analysis, we re-sampled the control group multiple times without considering the stratified group and compared the IgG(S) titers and T-spot counts with the breakthrough infection group and obtained similar results.

Taken together, our findings suggest that the serum IgA(S) responses in the early stages following booster vaccination may serve as a valid predictor of breakthrough infection risk, whereas a single measurement of IgG(S) antibody titers and T-spot counts may not.

## Discussion

To date, a total of 13.64 billion doses of COVID-19 vaccines have been administered worldwide. Approximately 70.7% of the global population has received at least 1 dose of COVID-19 vaccine. However, in low-income countries, this number drops to 32.8% because of global disparities in vaccine access. In 2022, the daily administration rate had significantly decreased from approximately 10 million to 100,000 doses, representing a staggering 99% reduction. By 2024, this figure had further declined to just 10,000 doses per day [48]. In the post-COVID-19 era, considering the high-level population immunity worldwide due to widespread infection and vaccination, the uptake of booster vaccines faces challenges, including vaccine hesitancy, decreasing vaccine coverage due to adverse reactions [49], and a declining interest in COVID-19 [50, 51]. In 2023, the WHO declared an end to the global COVID-19 emergency. However, both the WHO and the Centers for Disease Control and Prevention continue to emphasize the importance of vaccination and maintaining preparedness for potential future outbreaks. They suggest that healthy individuals aged 6 years and older may not necessarily require a booster dose, but individuals aged 65 years and older or those who are severely immunocompromised should receive an additional dose 6 months after their last vaccination [7, 52]. With new vaccination programs tailored for the post-COVID-19 era, distinct from the rollout strategies designed for strict pandemic control, it is crucial to further refine the prioritization of individuals for booster doses and to determine the optimal timing of these doses based on the risk of breakthrough infection. This approach seeks to manage COVID-19 effectively through a practical and cost-efficient vaccination strategy.

Previous studies generally investigated differences in vaccine-elicited antibody responses after stratification of the population according to variables including age, sex, lifestyle habits, comorbidities, adverse reactions, and medication use (i.e., in “given” groups) [28, 53–57]. Our high-throughput quantification of IgG(S) in over 2,500 participants in the Fukushima vaccination cohort revealed that individual-level antibody dynamics (i.e., the time-course patterns) elicited by primary and additional vaccinations can be stratified into 6 and 4 groups, respectively, by using a combined approach with process-based mathematical modeling and data-driven analysis (i.e., without determining these “given” groups in advance). Especially, we have identified 3 notable groups: the durable (G1 and g1), vulnerable (G6 and g4), and rapid-decliner (G4 and g3) populations. Additionally, extensive T-spot analyses conducted at multiple time points in 854 participants showed a potential correlation between the durability and vulnerability of booster vaccine-elicited antibody dynamics and cellular immune responses. Furthermore, an analysis of IgA(S) titers measured in 268 participants suggested that, unlike IgG(S) titers and T-spot counts, IgA(S) titers may predict the risk of breakthrough infection during the early stages after booster vaccination.

The results of previous studies, including our own [13, 58, 59], have established that individuals with immune-suppressive diseases have relatively low rates of seroconversion or generate low levels of SARS-CoV-2 antibodies after receiving primary and/or booster vaccinations, especially compared to healthy populations. This is particularly evident among organ transplant recipients. Because multiple booster vaccinations have demonstrated their effectiveness in enhancing both humoral and cellular immunity for the majority of individuals, including those who are immunocompromised and organ transplant recipients, it becomes imperative to prioritize these patients for additional booster doses. Additionally, our analysis indicates that certain populations, specifically the vulnerable (g4) and rapid-decliner (g3) populations, may not maintain their antibody titers at a high enough level for an extended period following booster vaccination. Indeed, we found that these 2 groups were at high risk for breakthrough infections following booster vaccination: g4 had a higher percentage of breakthrough infections compared to the other groups, while g3 experienced breakthrough infections more rapidly. Considering that neutralizing antibody titers strongly correlate with vaccine effectiveness against symptomatic and severe infections [23–26], targeted interventions for individuals in g3 and g4, such as administering additional booster doses or passive antibody therapy [60], may be warranted. Therefore, it will be important in the post-COVID-19 era, as well as during future pandemics, to evaluate the impact of additional booster doses on individuals predicted to be part of vulnerable or rapid-decliner populations.

Our findings indicate that predicting the risk of breakthrough infection using only single measurements of IgG(S) titers and T-spot counts taken in the early phase after booster vaccination (an average of 53.6 days post-booster) is challenging, although the post-booster stratified groups correlated well with the risk. This difficulty may arise from the time-course patterns of IgG(S), particularly in the rapid-decliner group G3, which initially exhibits high IgG antibody levels and T-spot counts but ultimately faces a higher risk of breakthrough infection. In contrast, we found that IgA titers measured from the same samples were statistically significantly lower in participants who experienced breakthrough infections than in the control cases. This finding aligns with previous studies: Individuals with high levels of IgA(S) in nasal mucosa and saliva have been reported to have a reduced risk of breakthrough infection with SARS-CoV-2 [46, 47, 61], proving the protective effect of mucosal IgA(S) against infection. Multiple cohort studies have indicated that mRNA vaccination correlates with systemic and mucosal IgA responses in individuals without prior SARS-CoV-2 infection, albeit at significantly lower levels than in those previously infected [62–64]. The level of protection provided by IgA(S) against SARS-CoV-2 infection is currently not known, and it is not at all clear whether the induction of such low levels of IgA(S) contributes to protection against infection. This study examined the anti-spike antibody responses induced by mRNA vaccination in confirmed SARS-CoV-2-naive participants, demonstrating that the serum IgA(S) levels induced by mRNA vaccination in individuals without prior SARS-CoV-2 infection correlated with a reduced risk of infection. Notably, not all individuals recovering from SARS-CoV-2 infection produce SARS-CoV-2-specific antibodies [65], and serological assays utilizing N-antibody have detected only around 80% of individuals with a prior history of SARS-CoV-2 infection [66]. Thus, our study cannot exclude the potential misidentification of previously infected individuals as those who were without prior infection. However, it has been established that these individuals in our study are seronegative for antibodies elicited by SARS-CoV-2 infection, and it is suggested that the magnitude of the infection-specific immune response is also limited, suggesting that serum IgA(S) levels in our study were potentially lower than those in previously infected individuals. Future research should further explore the relationship between IgA antibodies and the risk of breakthrough infection, aiming to enable earlier and more accurate predictions. Furthermore, the mechanism via which intramuscularly injected mRNA vaccines elicit systemic and mucosal IgA responses is not yet understood. This study indicates that even minimal serum IgA levels can partially protect against infections. Furthermore, elucidating the mechanism by which mRNA vaccines stimulate serum and mucosal IgA may yield critical insights that can lead to more effective vaccinations against respiratory viral infections.

There are other limitations to this study that we can improve for a better understanding of the variation in booster vaccine-elicited immune dynamics at the individual level. Firstly, our analysis only incorporated datasets from the primary and booster vaccinations. However, it is crucial to note that we observed numerous transitions between stratified groups resulting from these vaccinations (i.e., from G1-G6 to g1-g4). To accurately identify the vulnerable and rapid-decliner populations who would benefit from additional booster doses, information after the last booster vaccinations, including “bivalent” vaccination, is crucial. Therefore, further analysis of the multi-modal longitudinal dataset, which includes the second, third, and subsequent booster vaccinations in the Fukushima vaccination cohort, is necessary to gain a better understanding of how the stratification changes as additional doses received. These analyses may also reveal individuals who, despite receiving multiple additional vaccinations, do not effectively mount an immune response, even if they are not elderly or immunocompromised patients. Specifically, conducting immunological analyses on these individuals will offer unique insights for the development of next-generation vaccinations. Our ongoing cohort is an ideal environment in which to evaluate why immunity, including humoral and cellular responses of vulnerable and rapid-decliner individuals, cannot be effectively elicited or maintained after additional booster vaccinations. Secondly, in the analysis of breakthrough infections after booster vaccination, accuracy may be limited in participants infected shortly after vaccination (i.e., those infected between visit 3 and visit 4). This limitation arises because reconstructing the antibody dynamics for these individuals using mathematical models was challenging (see **Supplementary Fig 1** and **Results** for details). Thus, these individuals need to be excluded from the stratification analysis, resulting in missing information on the post-booster stratified group. Additionally, participants who did not report a specific infection date, regardless of infection timing, were excluded from the analysis (**Fig 4A**). These cases were often individuals unaware of their infection timing, such as those with asymptomatic infections. Moving forward, it will be necessary to conduct follow-up surveys to collect precise infection dates. This will enable more accurate insights into participants experiencing early breakthrough infections.

In conclusion, identifying individuals within high-risk populations who do not maintain a booster vaccine-elicited immune dynamics for an extended period is a crucial task in the post-COVID-19 era. Guiding these individuals towards receiving additional doses is essential, and our study aims to make a contribution towards this endeavor [67]. The approach established in our research not only offers insights into post-COVID-19 vaccination programs but also makes a significant contribution to the early establishment of vaccination programs during future pandemics caused by unknown infectious diseases [68].

## Methods

### Ethics statement

The study was approved by the ethics committees of Hirata Central Hospital (number 2021-0611-1) and Fukushima Medical University School of Medicine (number 2021-116). Written informed consent was obtained from all participants individually before the survey.

### Participant recruitment and blood sample collection

The study design has been previously reported [10]. Briefly, the candidates were mainly recruited from Hirata village, Soma city, and Minamisoma city in rural Fukushima prefecture. We conducted non-sequential blood sampling and sequential blood sampling. A total of 2,526 individuals participated in non-sequential blood sampling (**Fig 1B**). The first vaccine dose was administered between March 10 and September 17, 2021; the 2nd dose between March 31 and October 8, 2021; and the booster dose between December 3, 2021, and August 9, 2022. The median interval for the primary 2-dose vaccine series was 21 days. The median interval between the 2nd primary dose and the booster dose was 231 days.

A total of 2,526 individuals participated in the 1st blood sampling (visit 1) to measure IgG antibody titers between September 9 and October 7, 2021, and 20 of these 2,526 blood samples were used to measure the neutralizing antibody titer by the live virus. A total of 2,443 individuals participated in the 2nd blood sampling (visit 2) to measure IgG antibody titers between November 21 and December 25, 2021, and 20 of these 2,443 blood samples were used to measure the neutralizing antibody titer by the live virus. A total of 2,363 individuals participated in the 3rd blood sampling (visit 3) to measure IgG antibody titers between February 24 and April 19, 2022, and 1,120, 20 and 268 of these 2,362 blood samples were used to measure IFN-γ, the neutralizing antibody titer by the live virus, and IgA antibody titers, respectively. A total of 2,250 individuals participated in the 4th blood sampling (visit 4) to measure IgG antibody titers between April 10 and July 29, 2022, and 1,795 and 164 of these 2,250 blood samples were used to measure IFN-γ and IgA antibody titers, respectively. A total of 970 individuals who did not receive a 2nd booster dose (i.e., 4th dose) participated in the 5th blood sampling (visit 5) for the measurement of IgG antibody titers between September 1 and November 16, 2022.

Furthermore, we performed additional blood sampling for 226 health care workers in the Fukushima vaccination cohort to measure antibody titers between May 31 and June 6, 2021, before the 4 main blood samplings (i.e., visits 1 through 4) described above. Note that the 12 health care workers (described in **Supplementary Fig 14**) with sequential blood sampling were not included in the non-sequential population of 2,526 participants. All blood sampling was performed at the medical facilities.

Of these 2,526 participants, 105 individuals were excluded due to missing data. After parameter estimation was performed on the remaining 2,421 individuals, 270 were further excluded due to defective data, either a short sampling interval or a small difference in antibody titers between 2 measurements (see below for details). A total of 130 “pre-infection” participants (i.e., infections before visit 1) were analyzed separately (**Fig 1AB** and **Supplementary Fig 1B**). Thus, the remaining 2,021 participants were used for unsupervised clustering analysis. In the booster dose analysis of these 2,021 participants, 378 individuals were excluded for not taking the booster or for missing data After parameter estimation was performed on the remaining 1,643 individuals, 25 “post-infection” participants (i.e., infections between visit 2 and visit 3) were subjected to a separate analysis (**Fig 1AB** and **Supplementary Fig 1B**). Hence, the remaining 1,618 participants were used for unsupervised clustering analysis after the booster vaccination (**Supplementary Fig 1A**). Then, a total of 141 participants who were infected after receiving a booster vaccination (i.e., between visit 3 and visit 5) were included in the analysis of the risk of breakthrough infection (**Fig 4A** and **Supplementary Fig 1B**).

### Basic demographic and health information

Information on sex, age, weight, height, daily medication, medical history, date of vaccination (primary/booster), adverse reaction after vaccination, type of vaccination, smoking habits, and drinking habits was retrieved from the paper-based questionnaire (summarized in **Table 1**).

### SARS-CoV-2-specific antibody measurement

All serological assays were conducted at The University of Tokyo. Specific anti-spike IgG (i.e., IgG(S)) and neutralizing activity against the Wuhan strain were measured as the humoral immune status after COVID-19 vaccination. Specific anti-nucleocapsid IgG antibody titers (i.e., IgG(N)) were used to determine past COVID-19 infection status. Chemiluminescent immunoassay with iFlash 3000 (YHLO Biotech, Shenzhen, China) and iFlash-2019-nCoV series (YHLO Biotech, Shenzhen, China) reagents were used in the present study. The measurement range was 2-3500 AU/mL for IgG(S) and 4-800 AU/mL for neutralizing activity. For neutralizing activity, AU/mL×2.4 was used to convert to International Units (IU/mL); for IgG(S), AU/mL×1.0 was used to convert to binding antibody units (BAU/mL). The testing process was as per the official guideline. Quality checks were conducted every day before starting the measurement.

### SARS-CoV-2 Roche anti-nucleocapsid antibody analysis

Serum samples were heat-inactivated at 56 °C for 30 min before use. Antibody titers for nucleocapsid (N) were measured using Elecsys Anti-SARS-CoV-2 (Roche, Basel, Switzerland) immunoassay kits, according to the manufacturer’s instructions, as described previously [69]. A cutoff index (COI) of 1.0, as determined by the manufacturer, was used to determine the presence or absence of anti-N antibody levels. Since the COVID-19 vaccines approved in Japan are only spike-based vaccines, anti-N antibodies are induced by infection but not by vaccines, whereas anti-S antibodies are induced by both infection and vaccines.

### SARS-CoV-2 anti-spike IgA antibody measurement

Serum samples were heat-inactivated at 56 °C for 30 minutes before testing. Anti-spike receptor binding domain (RBD) IgA levels against the ancestral strain and against the BA.1, BA.2, BA.5, BQ.1, and XBB.1 variants in sera were measured using the V-PLEX SARS-CoV-2 Panel 33 kit (Meso Scale Discovery), following the manufacturer’s instructions. The minimum dilution was set at 1:1000. Samples with values below the detection limit were assigned half of the detection limit.

### Neutralizing antibody titer assay

According to our previous method, the viral neutralizing titers in the serum samples were determined as a 50% neutralizing titer (NT_50_) [70]. In brief, serum samples were serially diluted 3-fold in medium, with an initial dilution of 1:10 (final dilution range 1:21,870). The diluted serum was incubated with 700 TCID_50_ units of recombinant GFP viruses carrying the variant spike gene (B.1.1 or BA.5) for 1 hour at 37°C and was then added to VeroE6/TMPRSS2 cells [71] seeded in black 96-well clear-bottom plates (PerkinElmer, MA, USA). After 1 hour, the cells were washed once and cultured in the complete culture medium. After the plate was incubated at 37°C for 34-36 hours, GFP fluorescence intensity was measured by using a microplate reader (Infinite®200 PRO, TEKAN, Mannedorf, Switzerland). NT_50_ values were derived by nonlinear regression ([agonist] vs. response – four-parameter variable slope) using GraphPad Prism v9.3.1 (Boston, MA, USA).

### SARS-CoV-2-specific cellular immunity measurement

Peripheral blood specimens were collected to perform the T-SPOT.COVID test (Oxford Immunotec, Abingdon, Cambridge, UK), which is a standardized Enzyme-Linked Immunospot (ELISpot) assay for measuring interferon-gamma (IFN-γ) release. The processing and analysis of samples followed the manufacturer’s instructions, as described in our previous study [72].

Briefly, the blood was drawn into lithium heparin tubes and then shipped overnight in temperature-regulated containers to LSI Medience Corporation in Tokyo, Japan. Upon arrival at the laboratory, the T-Cell Xtend reagent (provided by Oxford Immunotec) was added, and peripheral blood mononuclear cells were isolated using density gradient centrifugation. The cells were subsequently washed, counted, and distributed at a density of 250,000 ± 50,000 cells per well in a 96-well plate. The antigen pool containing SARS-CoV-2 structural proteins was utilized to stimulate T-cells in vitro. Following ELISpot analysis with the S6 TATC Entry Analyzer (CTL Corporation, Cleveland, Ohio, USA), the spot counts were categorized into 3 groups: nonreactive (≤ 4 spots), borderline (5-7 spots), and reactive (> 8 spots). An invalid test was determined if the number of spots in the negative control exceeded 10.

### Screening of breakthrough infections after booster vaccination

We conducted a double screening to accurately identify participants who developed breakthrough infections after booster vaccination. First, we identified 148 participants with a first-time IgG(N) titer of ≥ 2 AU/mL at either visit 4 or visit 5 [73, 74]. We then performed the Roche diagnostic immunoassay on blood samples from these participants collected after they had received a booster vaccination. We confirmed that 141 participants had positive results for the anti-N antibody test at either visit 4 (51 participants) or visit 5 (90 participants), indicating that these individuals experienced breakthrough infections after booster vaccination (**Fig 4C**).

### Modeling vaccine-elicited antibody dynamics

In our previous study, we developed mathematical models describing COVID-19 vaccine-elicited antibody dynamics to evaluate the impact of the primary 2 doses and booster dose of COVID-19 vaccinations [10, 31]. Here we used the following mathematical model:

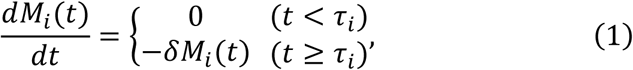

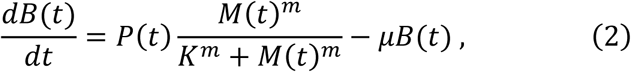

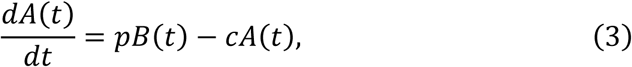

where the variables *M*_*i*_(*t*), *B*(*t*), and *A*(*t*) are the amount of mRNA inoculated by the *i*-th vaccination, the number of antibody-secreting cells, and the antibody titers at time *t*, respectively. The parameters *τ*_*i*_ and *δ* represent the timing of the *i*-th vaccination and the decay rate of mRNA, respectively. We denote *D*_*i*_ as the inoculated dose of mRNA by the *i*-th vaccination, that is, *M*_*i*_(*τ*_*i*_) = *D*_*i*_ and *M*_*i*_(*t*) = 0 for *t* < *τ*_*i*_. Because the data we used here were confined to time-course vaccine-elicited IgG(S) titers, one compartment of B cells including heterogeneous cell populations that produce antibodies (i.e., short-lived and long-lived antibody-secreting cells) was assumed. Thus, the average B cell population dynamics are derived in Eq.(2), where the product of *P*(*t*) and *M*(*t*)^*m*^/(*M*(*t*)^*m*^ + *K*^*m*^) represents the average *de novo* induction of the antibody-secreting cells. Here *P*(*t*) and *M*(*t*) are step functions that satisfy that *P*(*t*) = *P*_*i*_ and 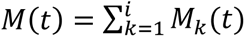 for *τ*_*i*_ + *η*_*i*_ ≤ *t* ≤ *τ*_*i*+1_ + *η*_*i*+1_ : otherwise *P*(*t*) = *M*(*t*) = 0, where *η*_*i*_ is the delay of induction of the antibody-secreting cells after the *i*-th vaccination, and *τ*_4_ = ∞. The parameters *m*, *K*, and *μ* correspond to the steepness at which the induction increases with increasing amount of mRNA (i.e., the Hill coefficient), the amount of mRNA satisfying *P*_*i*_/2, and the average decay rate of the antibody-secreting cell compartment, respectively. The other parameters, *p* and *c*, represent the antibody production rate and the clearance rate of antibodies, respectively.

Since the clearance rate of antibody is much larger than the decay of antibody-secreting cells (i.e., *c* ≫ *μ*), we made a quasi-steady state assumption, *dA*(*t*)⁄*dt* = 0, and replaced Eqs.(3) and (6) with *A*(*t*) = *pB*(*t*)⁄*c*. Moreover, since Eq.(1) is a linear differential equation, it can be rewritten as *M*_*i*_(*t*) = *D*_*i*_*e*^−*δt*^ for *t* ≥ *τ*_*i*_ : otherwise *M*_*i*_(*t*) = 0. Thus, Eqs.(1-3) are further simplified assuming *τ*_1_ = 0 and *D*_1_ > 0 as the following single ordinary differential equation:

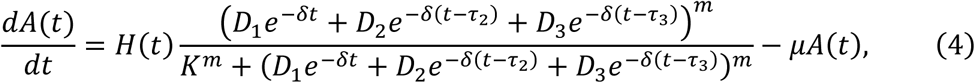

where *H*(*t*) = *H*_*i*_ = *pP*_*i*_/*c* for *τ*_*i*_ + *η*_*i*_ ≤ *t* < *τ*_*i*+1_ + *η*_*i*+1_, *τ*_4_ = ∞ and *D*_*i*_ > 0 for *τ*_*i*_ + *η*_*i*_ ≤ *t* : otherwise *D*_*i*_ = 0 (*i* = 1, 2, 3). This simple model can quantify the vaccine-elicited time-course antibody dynamics as described in **Supplementary Fig 2** under an arbitrary threshold of antibody titers *A*_TH_.

### Quantifying vaccine-elicited time-course antibody dynamics

In addition to the participants in the cohort, we included 12 health care workers whose serum was sequentially sampled for 340 days (on average 37 samples per individual) for validation and parameterization of the mathematical model for vaccine-elicited antibody dynamics: their serum was sampled for 260 days (on average 29 samples per individual) and 80 days (on average 8 samples per individual) after the primary and booster vaccinations, respectively. A nonlinear mixed effects model was used to fit the antibody dynamics model, given by Eq.(4), to the longitudinal antibody titers of IgG(S) obtained from the 12 health care workers. The mathematical model included both a fixed effect and a random effect in each parameter. That is, the parameters for individual *k*, *θ*_*k*_(= *θ* × *e*^*πk*^), are represented as a product of *θ* (a fixed effect) and *e*^*πk*^ (a random effect). The random effect *π*_*k*_ follows a Gaussian distribution with mean 0 and standard deviation Ω. Here we used lognormal distributions as prior distributions of each parameter to guarantee positiveness (negative values do not biologically make sense).

In the parameter estimation for antibody dynamics induced by the primary 2 vaccinations, we adapted the same method as used in our previous study [10]. That is, introducing parameters *f*_delay_ and *f*_degree_, we put *η*_1_ = *η*_*p*_, *η*_2_ = *f*_delay_*η*_*p*_, *H*_2_ = *H*_*p*_ and *H*_1_ = *f*_degree_*H*_2_ since *η*_1_ > *η*_2_ and *P*_1_ < *P*_2_. Then we estimated *η*_*p*_, *H*_*p*_, 0 < *f*_delay_ < 1 and 0 < *f*_degree_ < 1 for conducting biologically reasonable estimations. Also, we denote *η*_3_ = *η*_*b*_ and *H*_3_ = *H*_*b*_. We here assumed that the parameters *η*_*p*_, *η*_*b*_, *H*_*p*_, *H*_*b*_, *f*_delay_, *f*_degree_ and *m* varied across individuals, whereas we did not consider interindividual variability in other parameters to ensure parameter identifiability. In addition, our previous study [10] implied that *m* and *H* are key parameters determining vaccine-elicited antibody dynamics. Thus, for more accurate comparison between antibody dynamics before and after booster vaccination, we assumed that *m* differs between the primary and booster vaccinations, that is, *m*(*t*) = *m*_*b*_ for *t* ≥ *τ*_3_ + *η*_3_: otherwise *m*(*t*) = *m*_*p*_. Note that the half-life of mRNA (i.e., log 2 /*δ*) and dose of mRNA (i.e., *D*_*i*_) are assumed to be 1 day [75] and 100 (*μg*/0.5mL) [76], respectively, in our model.

Fixed effect and random effect were estimated by using the stochastic approximation expectation-approximation algorithm and empirical Bayes’ method, respectively. Fitting was performed using MONOLIX 2021R2 (www.lixoft.com) [77]. The estimated (fixed and individual) parameters are listed in **Supplementary Table 1**. The reconstructed individual-level time-course antibody dynamics calculated using estimated parameters are shown in **Supplementary Fig 14**. We found that most of the best-fitted estimated parameters in the mathematical model (i.e., *μ*, *K*, *η*, *f*_delay_, *f*_degree_) were the same or similar across the 12 individuals compared with those of parameters of *m*_*p*_, *m*_*b*_, *H*_*p*_ and *H*_*b*_ (see **Supplementary Table 1**). We note that (*m*_*p*_, *H*_*p*_) and (*m*_*b*_, *H*_*b*_) which showed wide variation of estimated values, contributed mainly to the vaccine-elicited antibody dynamics after the 2nd dose of primary and booster vaccinations, respectively, whereas the other parameters contributed to that after the 1st dose. In fact, what we are interested in is large variations after the primary and booster vaccinations (but not the negligible variation after the 1st dose). Therefore, we hereafter fixed the parameters in our mathematical model to be the estimated population parameters listed in **Supplementary Table 1**, except *m*_*p*_, *m*_*b*_, *H*_*p*_ and *H*_*b*_. These assumptions enabled us to accurately reconstruct the large variations in antibody dynamics after the primary and booster vaccinations, and these 4 parameters were independently estimated from each IgG(S) by a nonlinear least-squares method, even if each participant in the Fukushima vaccination cohort had only 4 or 5 measurements of antibody titers at different time points. Using the estimated parameters for each participant, we fully reconstructed the dynamics of IgG(S) titers of all participants after the 1st vaccination in **Supplementary Fig 2A**. The best-fit antibody titer curves of 100 randomly selected individuals are plotted along with the observed data for visualization in **Supplementary Fig 15**.

### Mathematical model validation

Outside the study period described in **Fig 1BCE**, to validate the reconstructed best-fit antibody titer curves described in **Supplementary Fig 2BC**, we prepared an additional 3rd or 4th antibody measurement of the Fukushima vaccination cohort around 3 to 4 months after the 2nd or 3rd timepoint data after the primary and booster vaccinations (**Supplementary Fig 1A**). Additional measurements were taken from 113 of 2,021 participants and 536 of 1,618 participants after the primary and booster vaccinations, respectively, and none of these participants were infected with COVID-19 (i.e., IgG(N)-negative participants) during that period. Then, comparing the reconstructed antibody titer curves and the additional data, we confirmed that the additional data points matched the prediction of our mathematical model with high accuracy (**Supplementary Fig 2BC**). Thus, our additional data and quantitative analysis demonstrated that the reconstructed best-fit antibody titer curves using our mathematical model are validated for the purposes of predicting antibody dynamics in a generalized population.

### Stratifying time-course pattern of antibody dynamics

As we previously assessed in [10, 31], we extracted the “features” described in **Supplementary Fig 7A** for each individual: the peaks (Peak^P^ and Peak^B^), durations (Duration^P^ and Duration^B^), and area under the curves (AUC^P^ and AUC^B^) of the reconstructed antibody dynamics before and after the booster vaccination, respectively. To quantify these features, we here assumed *A*_TH_= 100 and determined 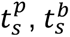 and 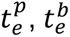 to correspond to the time for the antibody titer to be greater than and smaller than *A*_TH_, respectively. Therefore, the durations and AUCs of the antibody titer are formulated by 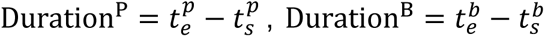 and 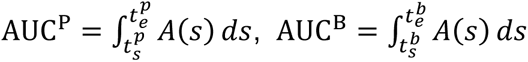, respectively. For 86 individuals, Peak^P^ was below *A*_TH_, and Duration^P^ and AUC^P^ were both determined to be 0. In **Supplementary Fig 7B**, we summarized distributions of the AUCs, durations, and peaks for all participants. Note that a similar trend was obtained under different *A*_TH_. In summary, we calculated a total of 6 features for each individual, in addition to our estimated individual parameters (i.e., *m*_*p*_, *H*_*p*_, *m*_*b*_, *H*_*b*_), which mainly contributed to the variation of vaccine-induced antibody dynamics. These features include 5 features for the primary vaccine-elicited antibody dynamics [Peak^P^, Duration^P^, AUC^P^, *m*_*p*_, *H*_*p*_], and 5 features for the booster vaccine-elicited antibody dynamics [Peak^B^, Duration^B^, AUC^B^, *m*_*b*_, *H*_*b*_], respectively.

In order to stratify the individual antibody response elicited by the primary and booster vaccinations, we used unsupervised random forest clustering based on each of 5 features. For the reconstructed antibody titer induced by the primary 2 vaccinations, the clustering method initially identified 13 distinct clusters (i.e., C1 to C13) (**Supplementary Fig 3AB**). First, we moved C6 and C10 to groups “unclassified” from further evaluation (270 individuals, around 10% of the 2,291 participants in the Fukushima vaccine cohort who were used in the stratification analysis for primary vaccination) on the grounds that their reconstructed antibody dynamics with estimated parameters may not be reasonable. This is because their antibody titers measured from 2 blood samples showed statistically significantly small differences (*p* < 1.0 × 10^−16^ by Welch’s two-sample t-test) and their sampling intervals were significantly shorter than the others (*p* = 0.01 by Welch’s two-sample t-test) (**Supplementary Fig 3C**).

Next, we amalgamated clusters exhibiting similar time-course patterns of antibody titer, and close distance on two-dimensional UMAP embeddings, resulting in a total 6 groups (i.e., G1 to G6): C8 to G1; C11 to G2; C1, C4, C12, and C13 to G3; C5 to G4; C7 and C9 to G5; and finally, C2 and C3 to G6 (**Fig 2A** and **Supplementary Fig 3D**). In contrast, for the reconstructed antibody titer induced by the booster vaccination, the clustering method initially identified 4 distinct clusters. Given that these 4 clusters clearly discriminated the time-course of antibody titer, we decided to use them as stratified groups (i.e., g1 to g4) (**Fig 2E** and **Supplementary Fig 3F**). We note that stratification of antibody dynamics by hierarchical clustering or decision trees was also performed, and this confirmed that our random forest provides a similar stratification compared to these conventional methods (see **Supplementary Fig 16** and **Supplementary Fig 17**).

### Selection of a control group for IgA antibody measurements

Among the 141 participants in the Fukushima vaccination cohort who experienced breakthrough infections after receiving a booster vaccination, IgA antibodies were measured retrospectively in 134 participants at visit 3 or visit 4 for whom serum samples were still available. To compare IgA titers between the breakthrough infection group and an uninfected control group, each breakthrough case was matched with uninfected controls based on gender, pre- and post-booster stratification groups, presence of underlying conditions, and vaccine type. From this pool, the control participant most closely matched in age and number of days to the 1st serum sample collection after the booster vaccination (i.e., days from date of booster dose to visit 3) was chosen. Specifically, if the age difference was within 5 years, the score was set to 0, with 1 point added for every additional 5-year increment. Similarly, if the difference in days to the 1st serum collection was within 10 days, the score was set to 0, with 1 point added for every 10-day increment beyond that. Control participants were randomly selected from those with the lowest total scores for each breakthrough infection case. The distribution of scores for the 134 control group participants was as follows: 56 had 0 points, 46 had 1 point, 21 had 2 points, 10 had 3 points, and 1 had 6 points.

### Statistical analysis

When necessary, the same variables were compared among different groups using Pearson’s chi-square test (for categorical variables), analysis of variance (ANOVA, for more than 2 numerical variables), Welch’s two-sample t-test, or pairwise Mann-Whitney U test (for 2 numerical variables). A Bonferroni correction was applied for multiple comparisons. All statistical analyses were performed using R (version 4.3.0).

## List of supplementary material

**Supplementary Figure 1** | The flowchart for all analysis

**Supplementary Figure 2** | Reconstructed dynamics of antibody titers at the individual level and validation

**Supplementary Figure 3** | Clustering of vaccine-elicited antibody response

**Supplementary Figure 4** | Quantifying fold increase of antibody titer by booster vaccination

**Supplementary Figure 5** | Longevity of primary and booster vaccine-elicited antibody titer

**Supplementary Figure 6** | Transition among stratified groups due to primary and booster vaccinations

**Supplementary Figure 7** | Quantifying of vaccine-elicited antibody dynamics

**Supplementary Figure 8** | Influence of infection history on vaccine-elicited antibody dynamics

**Supplementary Figure 9** | Correlation between T-spot counts and features of antibody dynamics

**Supplementary Figure 10** | Survival analysis of breakthrough infection for all participants who reported an infection date

**Supplementary Figure 11** | Correlation between IgA titers measured in nasal and serum samples

**Supplementary Figure 12** | Correlation between IgA(S) titers to the ancestral strain and to other strains

**Supplementary Figure 13** | Correlation between IgA titers, IgG titers, and T-spot counts

**Supplementary Figure 14** | Calibrating primary and booster vaccine-elicited antibody dynamics

**Supplementary Figure 15** | Reconstructed antibody titer trajectory for individual participants in Fukushima vaccination cohort

**Supplementary Figure 16** | Unsupervised stratification of vaccine-elicited antibody response with hierarchical clustering

**Supplementary Figure 17** | Unsupervised stratification of vaccine-elicited antibody response with a decision tree

**Supplementary Table 1** | Estimated fixed and individual parameters for 12 health care workers

## Acknowledgments

We would like to thank all the staff from Fukushima Medical University, Seireikai health care group, Hirata Village office, Soma City office, Soma Central Hospital, Soma General Hospital, Minamisoma City office, Minamisoma City Medical Association, Minamisoma Municipal General Hospital, Shindo Clinic Medical Governance Institute, and National Institute of Infectious Diseases, who contributed significantly to the accomplishment of this research, especially, Ms. Yuka Harada, Ms. Serina Noji, Ms. Naomi Ito, Dr. Makoto Kosaka, Mr. Anju Murayama, Mr. Sota Sugiura, Mr. Manato Tanaka, Ms. Yuna Uchi, Mr. Yudai Kaneda, Mr. Masahiko Nihei, Mr. Hideo Sato, Ms. Rie Yanai, Ms. Yasuko Suzuki, Ms. Keiko Abe, Dr. Hidekiyo Tachiya, Mr. Kouki Nakatsuka, Dr. Ryuzaburo Shineha, Ms. Miki Sato, Dr. Masahiko Sato, Mr. Naoharu Tadano, Mr. Kazuo Momma, Mr. Shu-ichi Mori, Ms. Saori Yoshisato, Ms. Katsuko Onoda, Mr. Satoshi Kowata, Mr. Masatsugu Tanaki, Dr. Tomoyoshi Oikawa, Dr. Joji Shindo, Ms. Xujin Zhu, Ms. Asaka Saito, Ms. Yuumi Kondo, Ms. Tomoyo Nishimura, and Ms. Emi Taeda. This study was supported by Medical & Biological Laboratories Co., Ltd., Shenzhen YHLO Biotech Co., Ltd., the distributor and manufacturer of the antibody measurement system (iFlash 3000), Research Center for Advanced Science and Technology in the University of Tokyo, and in part by Scientific Research (KAKENHI) B 23H03497 (to S.I.); Grant-in-Aid for Transformative Research Areas 22H05215 (to S.I.); Grant-in-Aid for Challenging Research (Exploratory) 22K19829 (to S.I.); AMED CREST 19gm1310002 (to S.I.); AMED Development of Vaccines for the Novel Coronavirus Disease, 21nf0101638 (to M.T. and S.I.); AMED Research Program on Emerging and Re-emerging Infectious Diseases 22fk0108509 (to S.I.), 23fk0108684 (to S.I.), 23fk0108685 (to S.I.); AMED Research Program on HIV/AIDS 22fk0410052 (to S.I.); AMED Program for Basic and Clinical Research on Hepatitis 22fk0210094 (to S.I.); AMED Program on the Innovative Development and the Application of New Drugs for Hepatitis B 22fk0310504h0501 (to S.I.); JST MIRAI JPMJMI22G1 (to S.I.); Moonshot R&D JPMJMS2021 (to K.A. and S.I.) and JPMJMS2025 (to S.I.); JST PRESTO JPMJPR19H5 (to Y.S.); JST CREST JPMJCR19H4 (to Y.S.); The National Research Foundation of Korea (NRF) grant funded by the Korea government (MSIT) (2022R1C1C2003637) (to K.S.K.); Shin-Nihon of Advanced Medical Research (to S.I.); SECOM Science and Technology Foundation (to S.I.); The Japan Prize Foundation (to S.I.); and Kowa Co (to M.T.).

## Author contributions

SI designed the research. MT conducted the data collection. HP, NN, KSK, KK, KA, and SI carried out the computational analysis. SI supervised the project. All authors contributed to writing the manuscript.

## Institutional review board statement

This study was approved by the ethics committees of Hirata Central Hospital (number 2021-0611-1) and Fukushima Medical University (number 2021-116). This study was conducted in accordance with the Code of Ethics of the World Medical Association (Declaration of Helsinki).

## Informed consent statement

Informed consent was obtained from all subjects involved in the study.

