## Supplementary information for "Longitudinal analysis of antibody titers after primary and booster mRNA COVID-19 vaccination can identify individuals at risk for breakthrough infection"

[1]interdisciplinary Biology Laboratory (iBLab), Division of Natural Science, Graduate School of Science, Nagoya University, Nagoya, Japan. [2]Department of Pathology, National Institute of Infectious Disease, Tokyo, Japan. [3]Department of Virology, Nagoya University Graduate School of Medicine, Nagoya, Japan. [4]Department of Scientific Computing, Pukyong National University, Busan, South Korea. [5]Department of Mathematics, Pusan National University, Busan, South Korea. [6]Department of Radiation Health Management, Fukushima Medical University School of Medicine, Fukushima, Japan. [7]Department of General Internal Medicine, Hirata Central Hospital, Fukushima, Japan. [8]Medical Governance Research Institute, Tokyo, Japan. [9]Department of Infectious Diseases, Nagoya University Hospital, Nagoya, Japan. [10]Department of Cardiology, Nagoya University Graduate School of Medicine, Nagoya, Japan. [11]Proteomics Laboratory, Isotope Science Center, The University of Tokyo, Tokyo, Japan. [12]Laboratory for Systems Biology and Medicine, Research Center for Advanced Science and Technology, The University of Tokyo, Tokyo, Japan. [13]Medical & Biological Laboratories Co., Ltd, Tokyo, Japan. [14]Department of Advanced Transdisciplinary Sciences, Hokkaido University, Sapporo, Hokkaido, Japan. [15]International Research Center for Neurointelligence, The University of Tokyo Institutes for Advanced Study, The University of Tokyo, Tokyo, Japan. [16]Department of Microbiology and Immunology, Faculty of Medicine, Hokkaido University, Sapporo, Japan. [17]Institute for Vaccine Research and Development (IVReD), Hokkaido University, Sapporo, Japan. [18]One Health Research Center, Hokkaido University, Sapporo, Japan. [19]AMED-CREST, Japan Agency for Medical Research

and Development (AMED), Tokyo, Japan. [20]Laboratory of Virus Control, Research Institute for Microbial Diseases, Osaka University, Suita, Japan. [21]Soma Medical Center of Vaccination for COVID-19, Fukushima, Japan. [22]Tokyo Foundation for Policy Research, Tokyo, Japan. [23]Institute of Mathematics for Industry, Kyushu University, Fukuoka, Japan. [24]Institute for the Advanced Study of Human Biology (ASHBi), Kyoto University, Kyoto, Japan. [25]Interdisciplinary Theoretical and Mathematical Sciences Program (iTHEMS), RIKEN, Saitama, Japan. [26]NEXT-Ganken Program, Japanese Foundation for Cancer Research (JFCR), Tokyo, Japan. [27]Science Groove Inc., Fukuoka, Japan. [28]Minamisoma Municipal General Hospital, Fukushima, Japan.

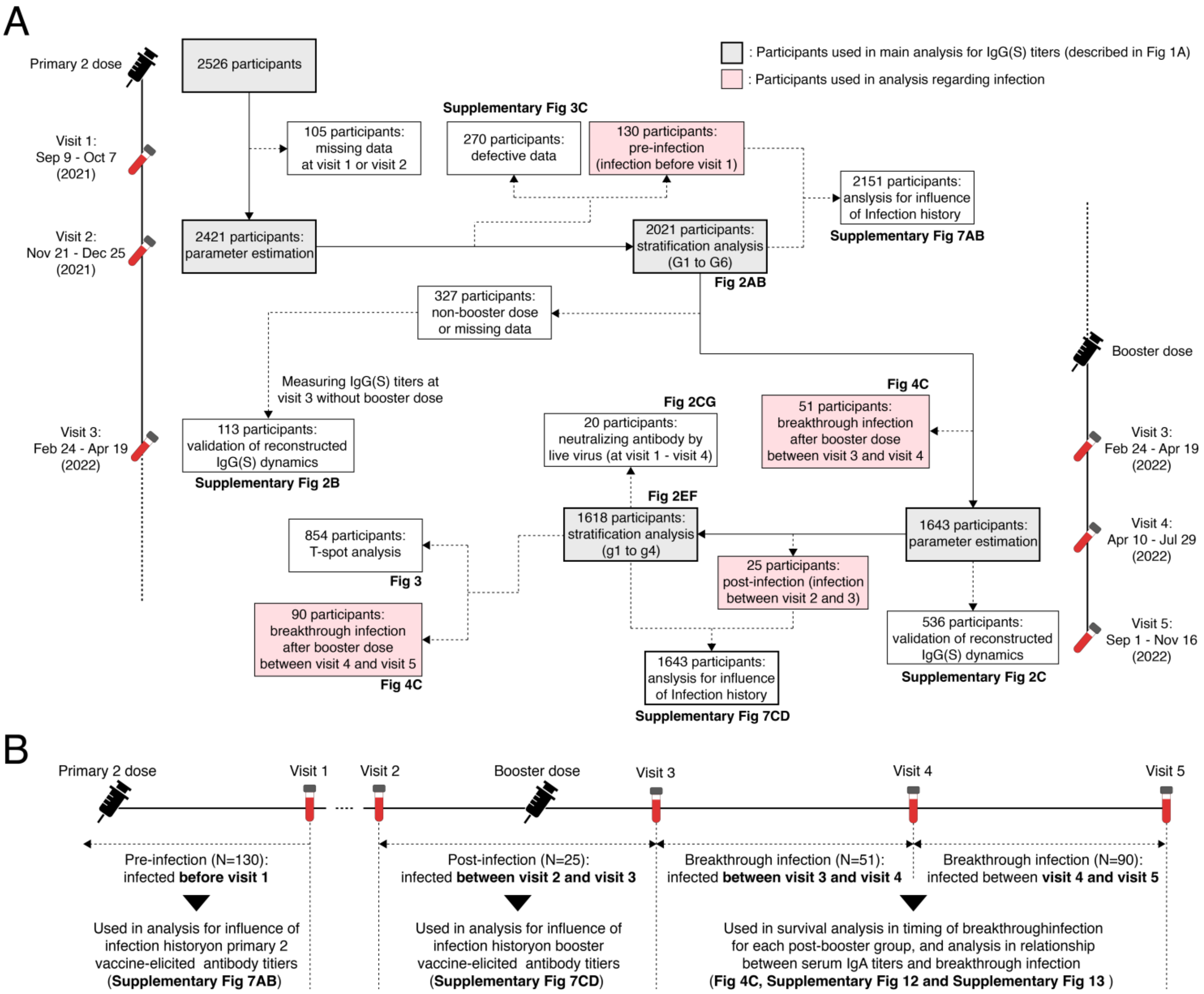


**Supplementary Figure 1. The flowchart for all analysis**: **(A)** The entire flowchart for the vaccination cohort is shown, detailing the number of participants and the inclusion criteria for our study. **(B)** This flowchart outlines the participants who developed infections during the study period. In particular, it describes the timing of infections categorized as pre-infection, post-infection, and breakthrough infection in our study, and the analyses conducted for participants in each infection group.

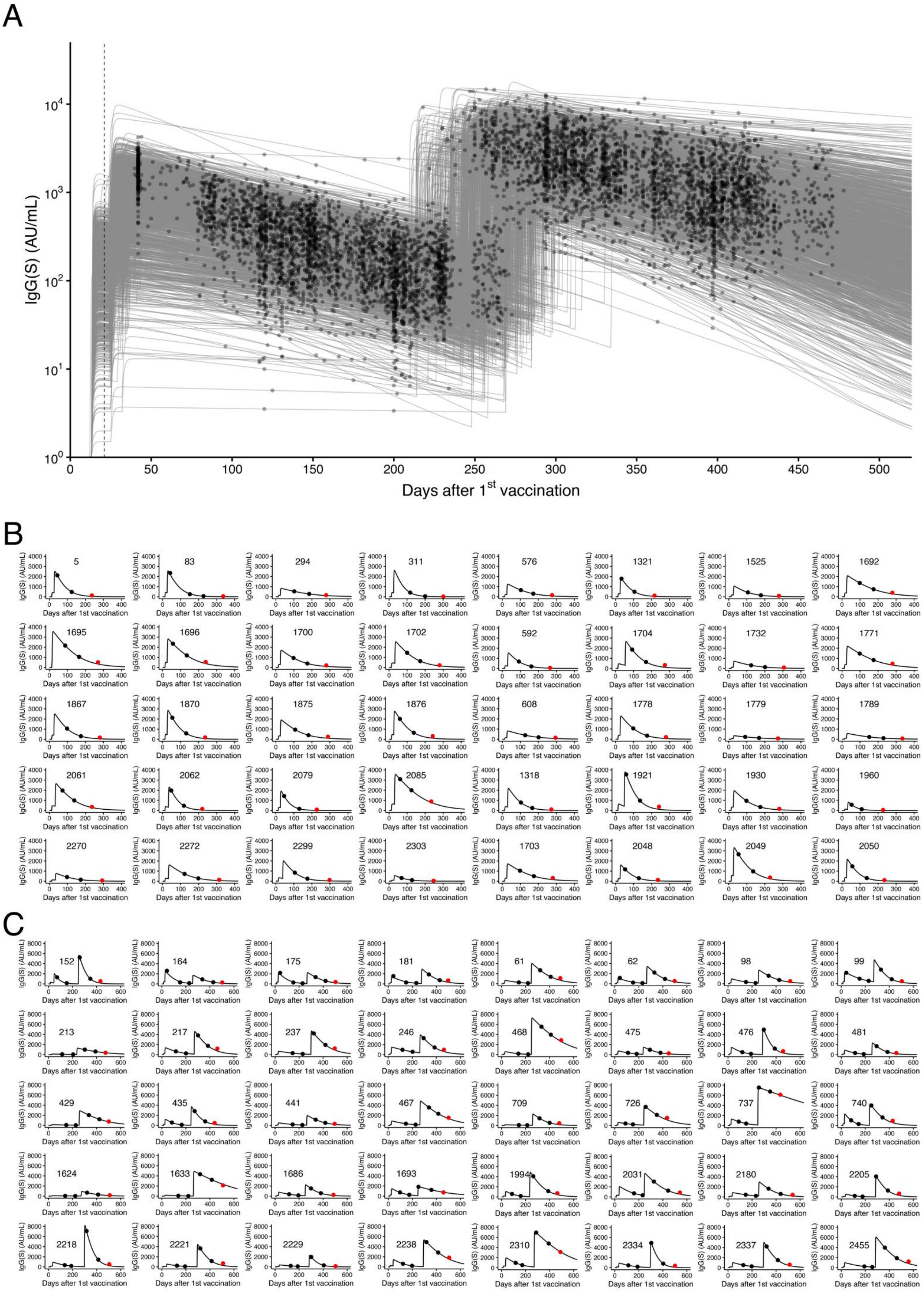
A
IgG(S) (AU/mL)
Days after 1st vaccination
B
5
83
294
311
576
1321
1525
1692
1695
1696
1700
1702
592
1704
1732
1771
1867
1870
1875
1876
608
1778
1779
1789
2061
2062
2079
2085
1318
1921
1930
1960
2270
2272
2299
2303
1703
2048
2049
2050
C
152
164
175
181
61
62
98
99
213
217
237
246
468
475
476
481
429
435
441
467
709
726
737
740
1624
1633
1686
1693
1994
2031
2180
2205
2218
2221
2229
2238
2310
2334
2337
2455

**Supplementary Figure 2. Reconstructed dynamics of antibody titers at the individual level and validation: (A)** Reconstructed time-course vaccine-elicited IgG(S) titers (gray curves) with observed IgG(S) titers (black circles) for all 1618 participants used in stratifying after booster vaccination are shown. **(B)(C)** Validation of the reconstructed IgG(S) titers of individual who did not receive the additional vaccination and did not get infected with COVID-19, for 40 randomly selected participants after **(B)** the primary and **(C)** booster vaccinations, respectively. The black curves describe the best-fit antibody titer curves. The black circles correspond to the observed IgG(S) titers used to estimate time-course IgG(S) titer, and the red circles correspond to the additionally observed IgG(S) titers.

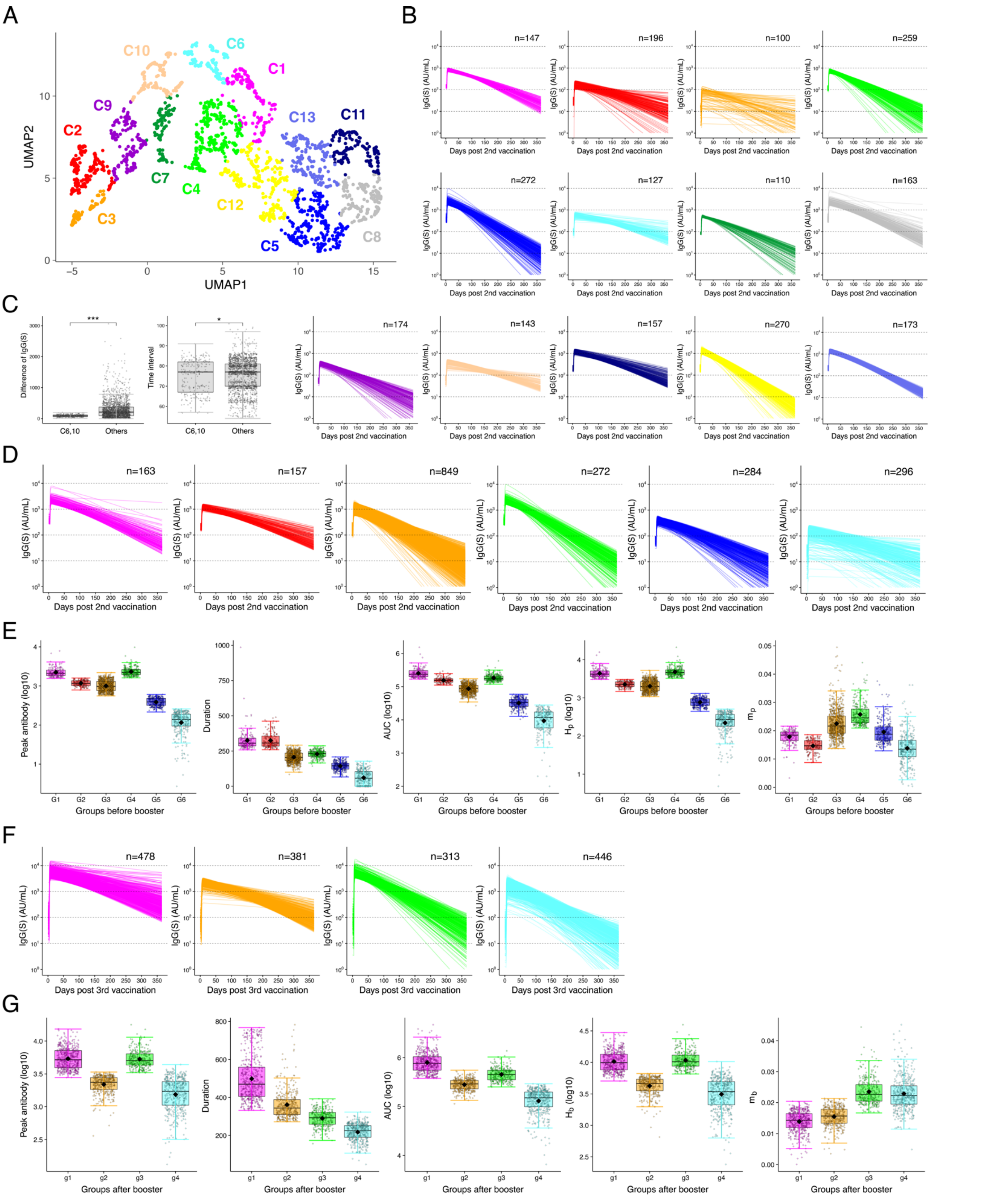


**Supplementary Figure 3. Clustering of vaccine-elicited antibody response: (A)** UMAP of 13 clustered antibody responses based on the extracted features from the reconstructed individual-level antibody dynamics are shown. Data points represent individual participants and are colored according to the 13 clusters (i.e., C1 to C13). **(B)** Reconstructed individual antibody dynamics after the first primary dose in each of the 13 clusters are shown in different colors. **(C)** The difference in antibody titers between the 2 measurements and its sampling interval are compared between C6, 10 and other clusters. **(D)** Reconstructed individual antibody dynamics in each of the 6 merged groups (i.e., G1 to G6) before booster vaccination are shown. **(E)** Distributions of 2 estimated parameters and 3 features extracted from the reconstructed antibody titers are shown for the 6 groups before booster vaccination. **(F)** Reconstructed individual antibody dynamics in each of the 4 post-booster groups (i.e., g1 to g4) are shown. **(G)** Distributions of 2 estimated parameters and 3 features extracted from the reconstructed antibody titers are shown for the 4 post-booster groups.

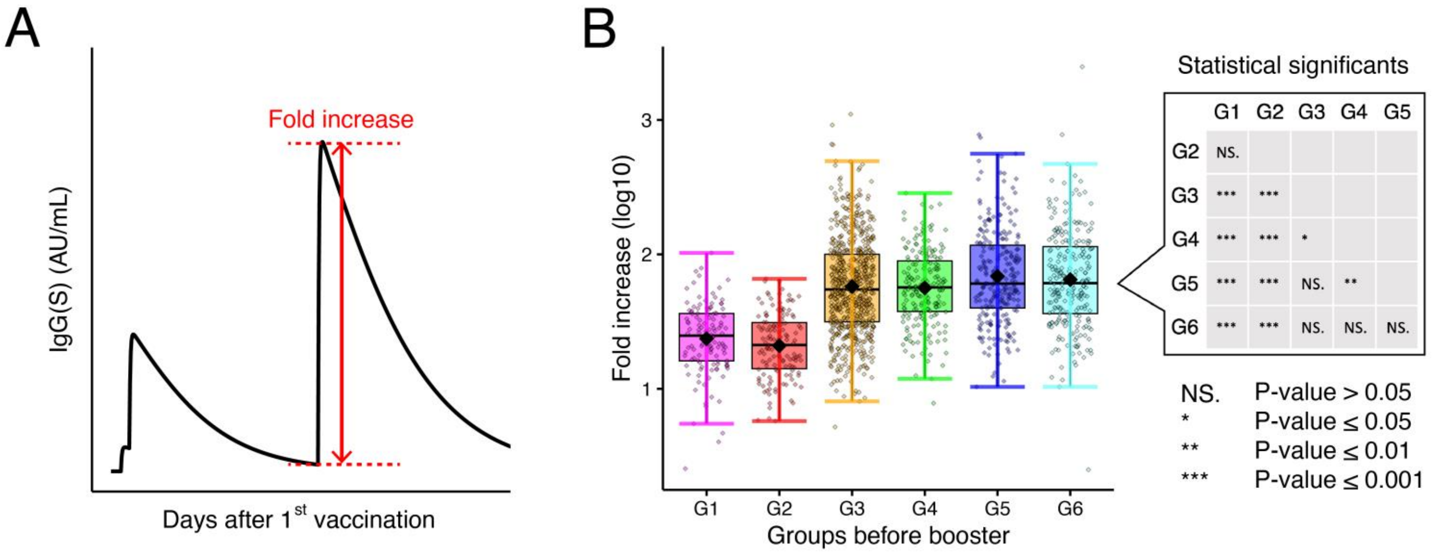


**Supplementary Figure 4. Quantifying fold increase of antibody titer by booster vaccination: (A)** The fold increase of antibody titer by the booster vaccination is defined as the difference between the minimum IgG(s) titer and the maximum titer around after the primary and booster vaccinations, respectively. **(B)** Distributions of the fold increase from the reconstructed individual-level antibody dynamics are plotted, respectively, among total or stratified groups G1 to G6.

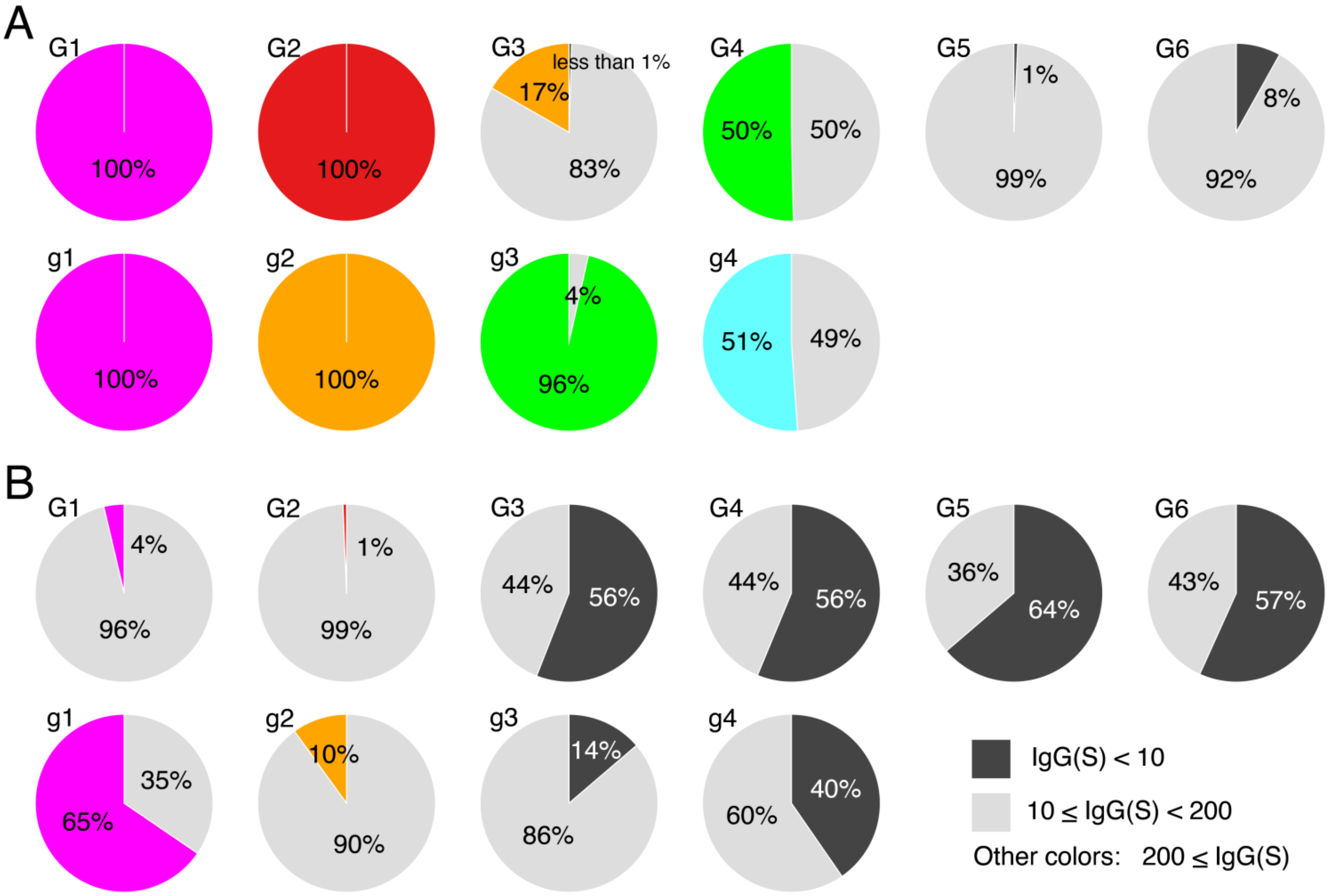


**Supplementary Figure 5. Longevity of primary and booster vaccine-elicited antibody titer:** Fraction of individuals with antibody titers <10, 10-200, and ≥200 AU/mL in each stratified group (top: G1 to G6, bottom: g1 to g4) at **(A)** 180 and **(B)** 365 days after the primary and booster vaccinations are plotted as black, white, and group color, respectively, in pie charts. Note that 10 AU/mL is a clear threshold for unvaccinated individuals [1], implying that individuals showing titers less than this threshold have little vaccine efficacy. In addition, 200 AU/mL is an indicator for another efficacy threshold because more than 80% of vaccinated persons maintained their antibody titers above 200 AU/mL for at most 3 months after the first vaccination regardless of group in our cohort.

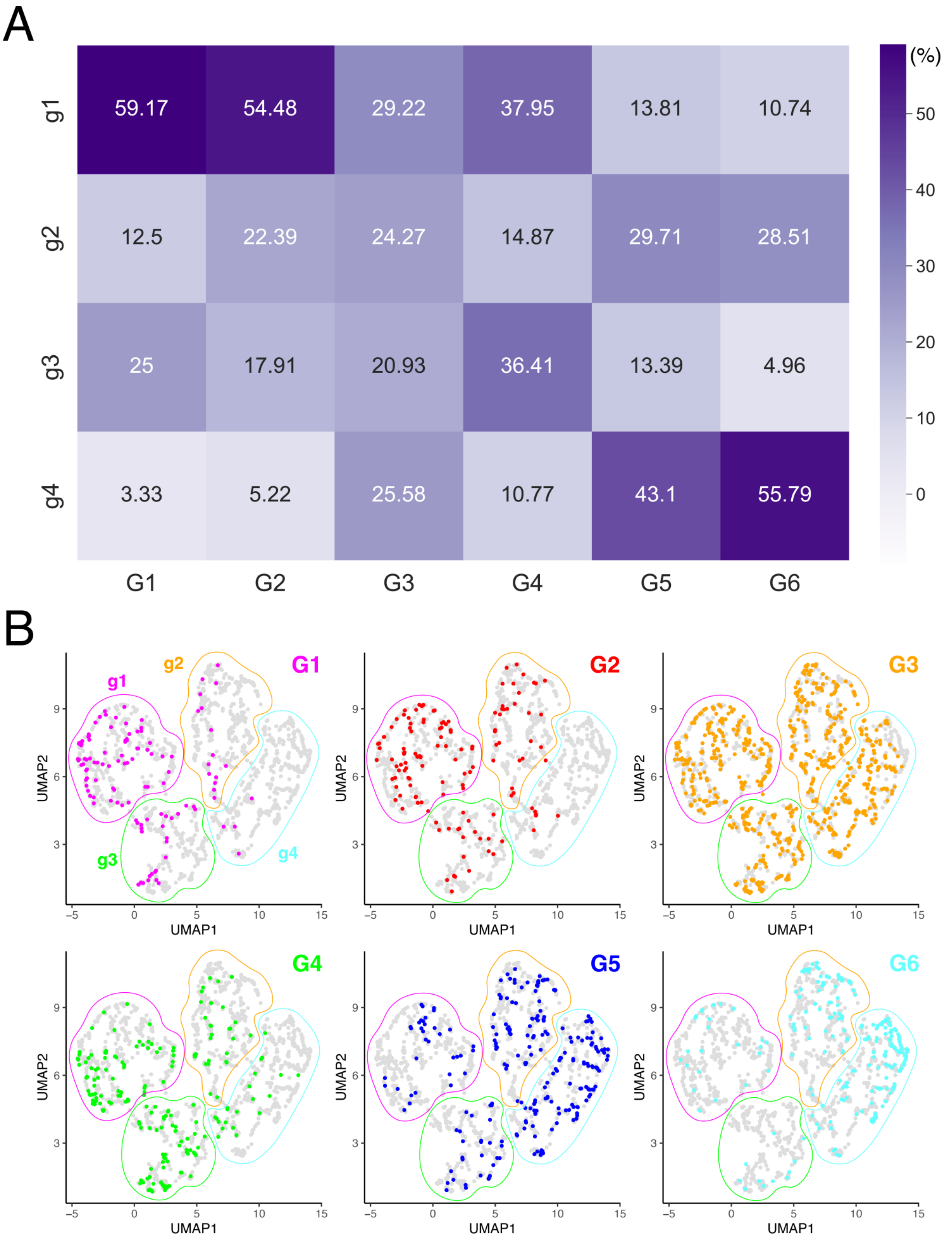

A
(%)
50
40
30
20
10
0
g1
g2
g3
g4
59.17
54.48
29.22
37.95
13.81
10.74
12.5
22.39
24.27
14.87
29.71
28.51
25
17.91
20.93
36.41
13.39
4.96
3.33
5.22
25.58
10.77
43.1
55.79
G1
G2
G3
G4
G5
G6
B
G1
G2
G3
G4
G5
G6
g1
g2
g3
g4
UMAP1
UMAP2

**Supplementary Figure 6. Transition among stratified groups due to primary and booster vaccinations: (A)** The transition matrix of participants from groups G1-G6 to groups g1-g4 is shown. The $ij$-element of the matrix ($i = 1, 2, \ldots, 6$, and $j = 1, 2, 3, 4$) represents the proportion of participants in G$i$ who transitioned to groups g$j$ after receiving the booster dose. **(B)** Each panel displays the UMAP shown in **Fig 2E**, with regions corresponding to g1 to g4 demarcated by colored borders (see top-left panel for a correspondence between the borders and groups g1 to g4). All participants are represented by gray dots on UMAP, but only participants belonging to G1, G2, ..., or G6 are marked with the color associated with their respective group.

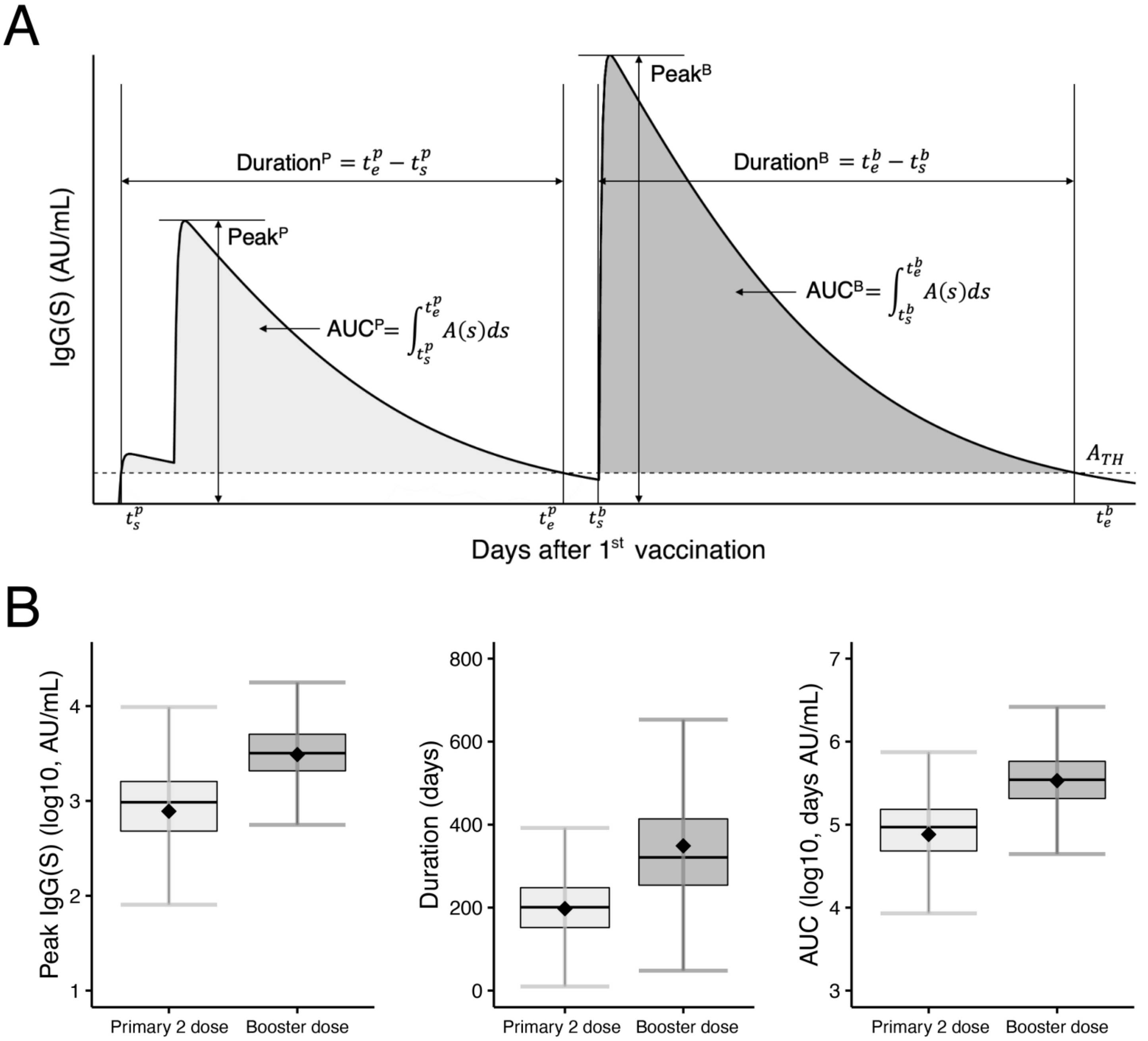


**Supplementary Figure 7. Quantifying of vaccine-elicited antibody dynamics: (A)** Vaccine-elicited antibody response after the first vaccination (i.e., $t = 0$) is described with the following "features": the peaks ($\mathrm{Peak}^{\mathrm{P}}$ and $\mathrm{Peak}^{\mathrm{B}}$), durations ($\mathrm{Duration}^{\mathrm{P}}$ and $\mathrm{Duration}^{\mathrm{B}}$), and area under the curves ($\mathrm{AUC}^{\mathrm{P}}$ and $\mathrm{AUC}^{\mathrm{B}}$) of the antibody titers elicited by the primary and booster vaccinations, respectively. The horizontal dashed line corresponds to the arbitrary threshold ($A_{\mathrm{TH}}$) for calculating the duration and AUC, respectively. **(B)** Distributions of the extracted features from the reconstructed antibody dynamics (i.e., the peak, duration, and AUC) for all participants are plotted. The dataset for each distribution was normalized by the value corresponding to the 95th percentile of data values, and data with values larger than this value were removed to improve the visibility of the figure.

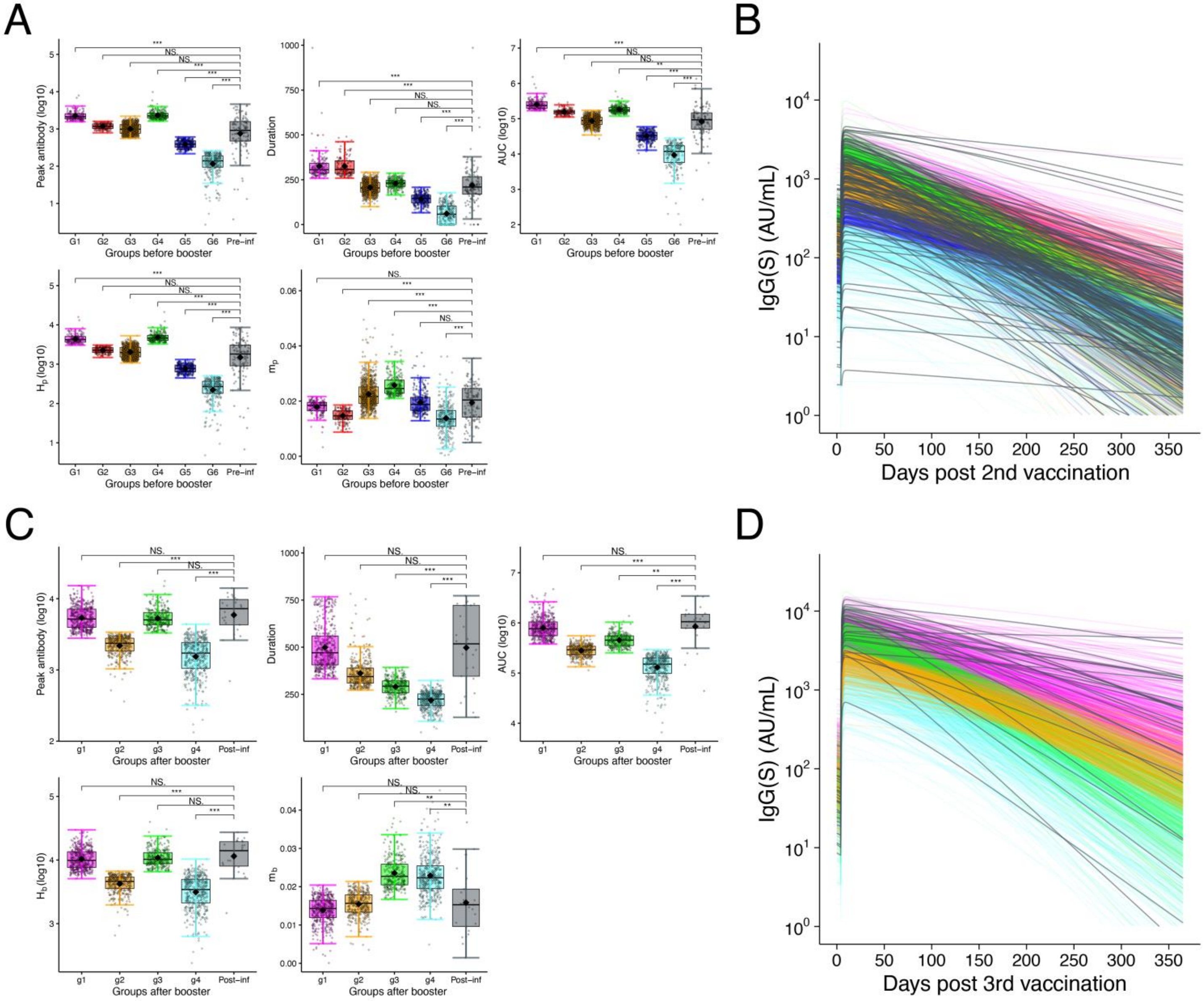


**Supplementary Figure 8. Influence of infection history on vaccine-elicited antibody dynamics: (A)(C)** The 3 features extracted from the reconstructed antibody titers and the 2 estimated parameters are compared between participants with and without infection history. **(A)** Comparison between 6 groups before booster vaccination (i.e., G1 to G6) and the pre-infection group, and **(C)** 4 groups after booster vaccination (i.e., g1 to g4) and the post-infection group, are shown, respectively. **(B)(D)** The reconstructed individual antibody dynamics for participants infected with COVID-19 before **(B)** the primary (pre-infection) and **(D)** the booster (post-infection) vaccinations, respectively, are described in black. The colored curves represent the reconstructed antibody dynamics for the stratified groups **(A)(B)** before and **(C)(D)** after the booster vaccination, which are presented in **Fig 2B** and **Fig 2F**, respectively. For **(A)** and **(C)**, statistical significances are calculated by the pairwise Mann-Whitney U test, and p-values are corrected by Bonferroni's method (NS.: p-value > 0.05, $*$: p-value $\leq$ 0.05, $**$: p-value $\leq$ 0.01, and $***$: p-value $\leq$ 0.001, respectively).

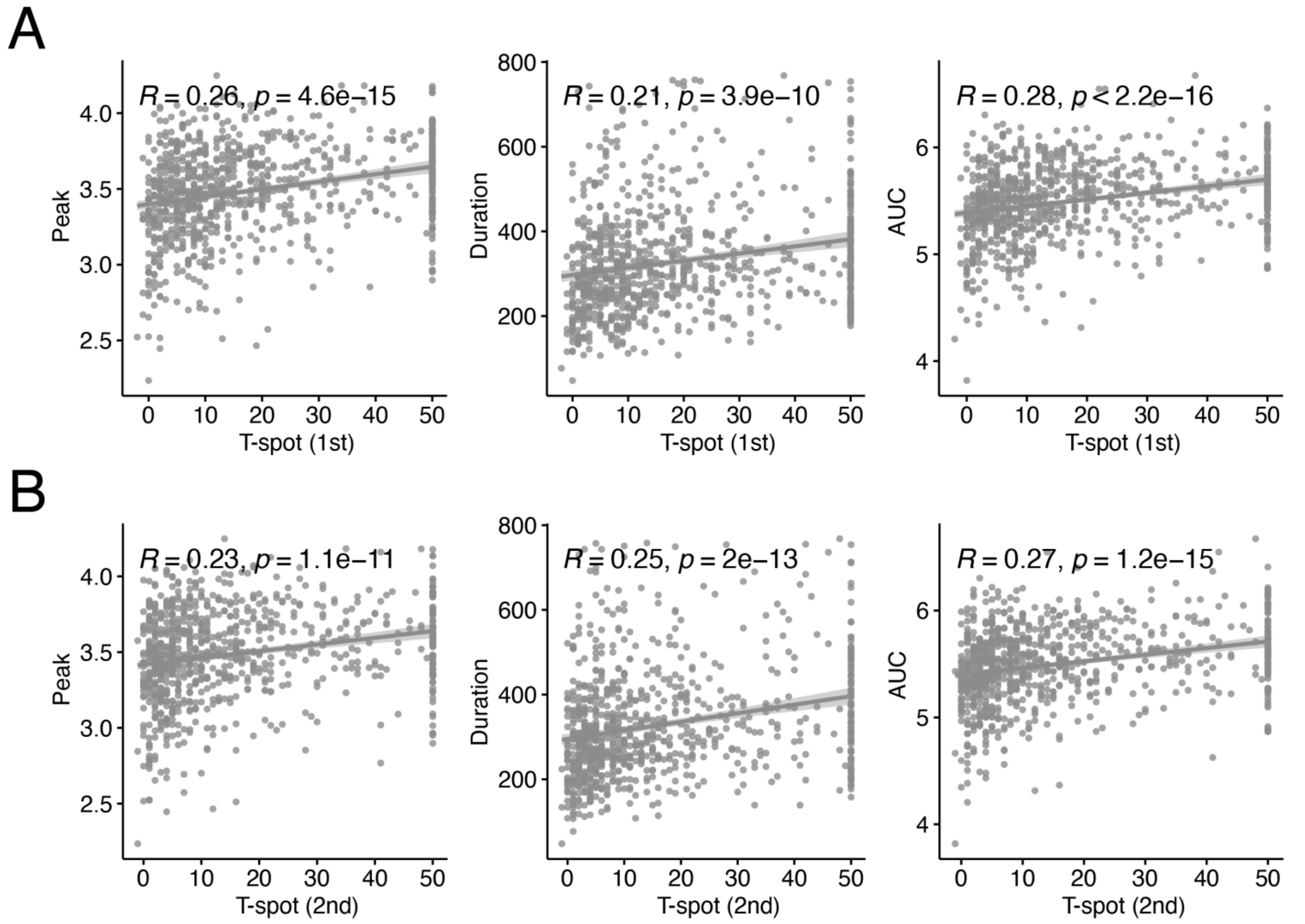


**Supplementary Figure 9. Correlation between T-spot counts and features of antibody dynamics:** The relationship between the **(A)** 1st and **(B)** 2nd measured T-spot response and the features of IgG(S) dynamics (peak, duration, and AUC) are shown, respectively. Regression lines with 95% CIs are shown, and the $R$ and $p$ on the top of each panel indicate the Pearson's correlation coefficient and its p-value, respectively.

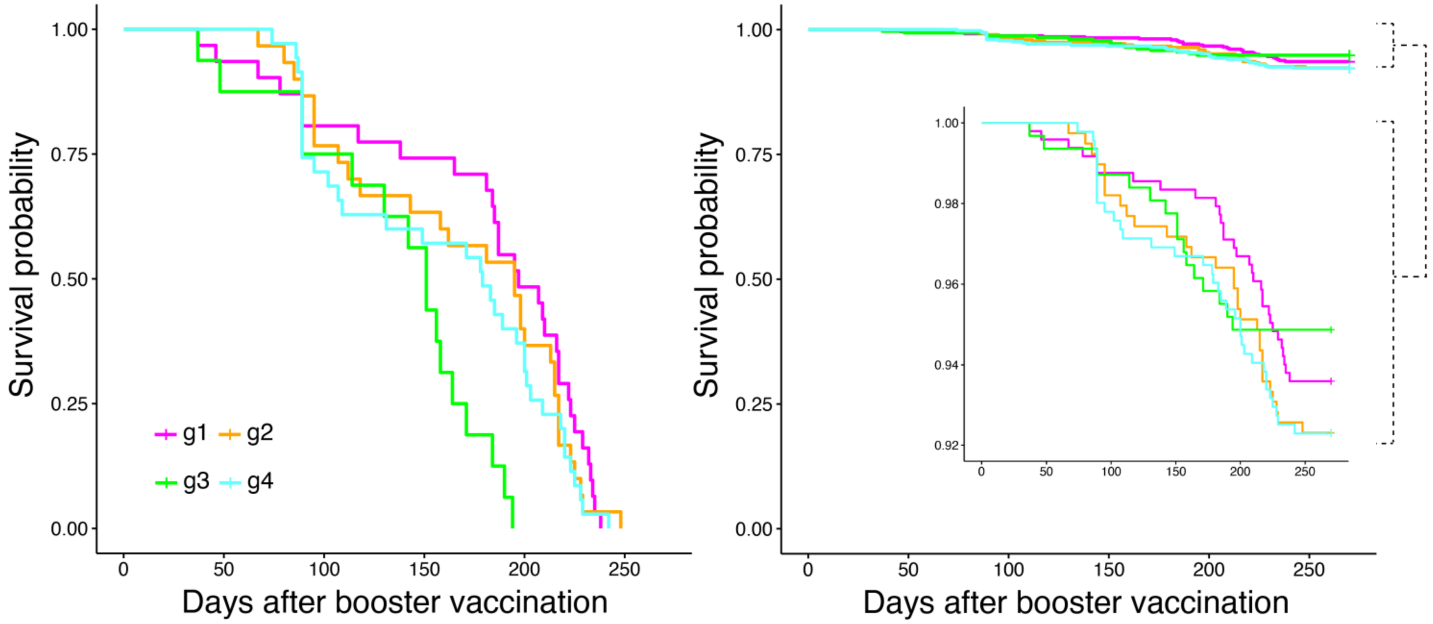


**Supplementary Figure 10. Survival analysis of breakthrough infection for all participants who reported an infection date:** The survival probability for each post-booster stratified group is presented for the participants with breakthrough infection only (left) and for the total population of each group (right). This analysis includes all 112 participants who experienced breakthrough infections after booster vaccination. Notably, in **Fig 4B**, 36 of the 112 participants were excluded from the post-booster stratification analysis because their IgG(S) dynamics could not be reconstructed using the mathematical model because they had only a single measured IgG(S) titer before infection. For these participants, their post-booster stratification group was estimated based on measured IgG(S) titers at visit 3: we assumed each participant belonged to the group with the smallest difference between their IgG(S) titer at visit 3 and the average curve of the g1-g4 groups (**Fig 2F**). Additionally, since these 36 participants reported only the month of infection and not specific dates, we estimated their infection date as the midpoint of the reported month. Consistent with the trends observed in **Fig 4B**, we found that the rapid-decliner group (g3) experienced breakthrough infections earlier than did the other groups. Furthermore, the vulnerable group (g4) demonstrated a higher frequency of breakthrough infections within the early stage after booster vaccination (within 100 days) compared to the other groups, whereas the durable group (g1) exhibited slower progression to breakthrough infection. Overall, the risk of breakthrough infection was higher in g3 and g4.

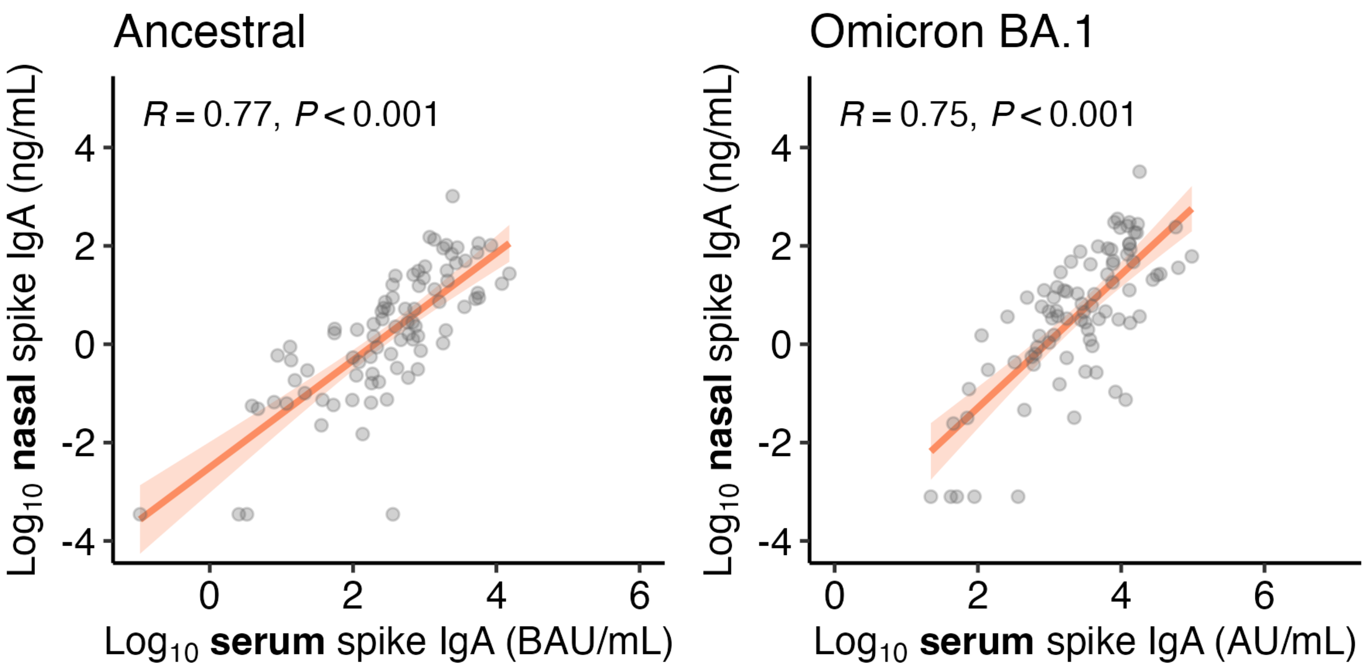


**Supplementary Figure 11. Correlation between IgA titers measured in nasal and serum samples:** The relationships between nasal and serum IgA(S) titers for symptomatic cases with matching collection dates against the ancestral strain (left) and BA.1 strain (right) are shown. Regression lines with 95% CIs are shown, and the $R$ and $P$ on the top of each panel indicate the Pearson correlation coefficients and their p-values, respectively.

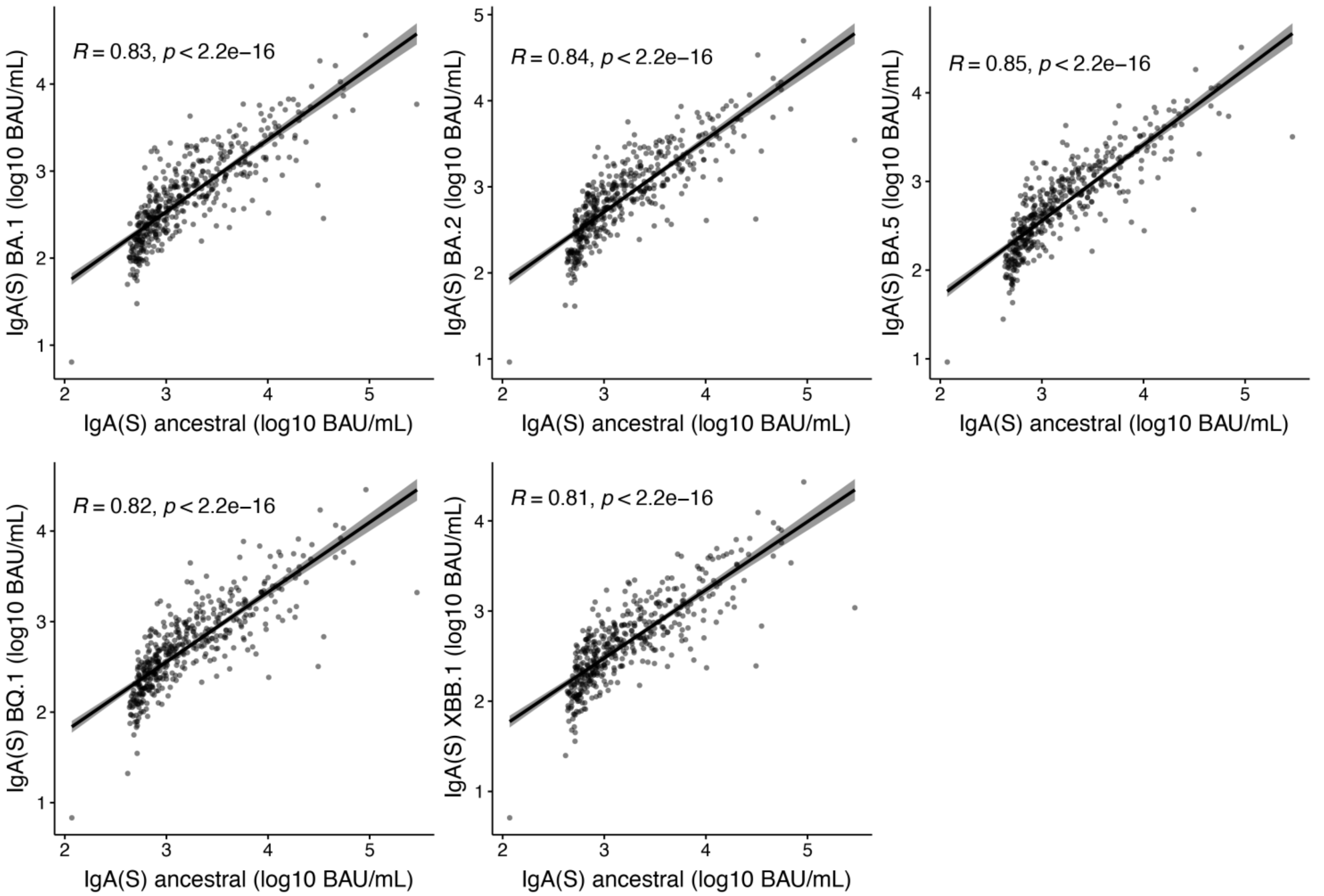


**Supplementary Figure 12. Correlation between IgA(S) titers to the ancestral strain and to other strains:** The relationships between IgA(S) titers against the ancestral strain, and IgA(S) titers against the BA.1, BA.2, BA.5, BQ.1, and XBB.1 strains are shown, respectively. Regression lines with 95% CIs are shown, and the $R$ and $p$ on the top of each panel indicate the Pearson correlation coefficients and their p-values, respectively.

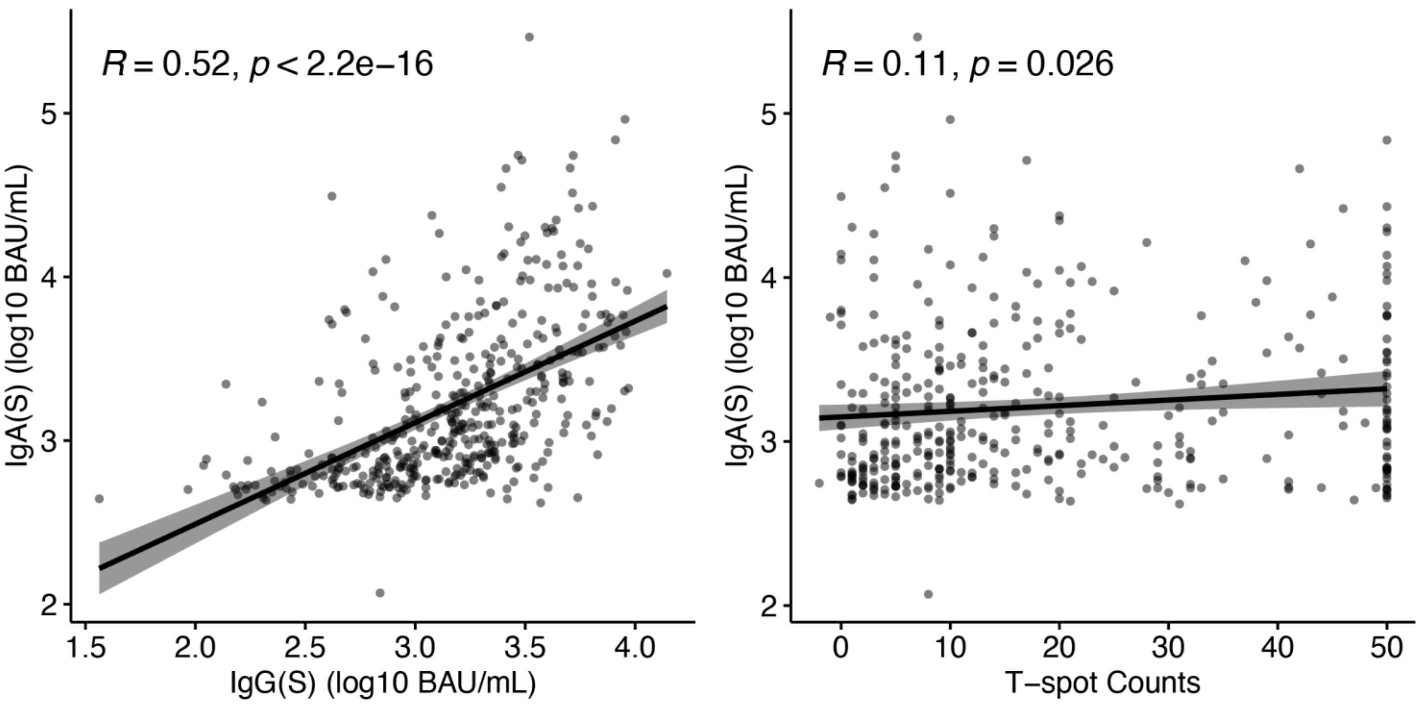


**Supplementary Figure 13. Correlation between IgA titers, IgG titers, and T-spot counts:** The relationships between IgA(S) titers and IgG(S) titers, as well as between IgA(S) titers and T-spot counts, are shown. Regression lines with 95% CIs are shown, and the $R$ and $p$ on the top of each panel indicate the Pearson correlation coefficients and their p-values, respectively.

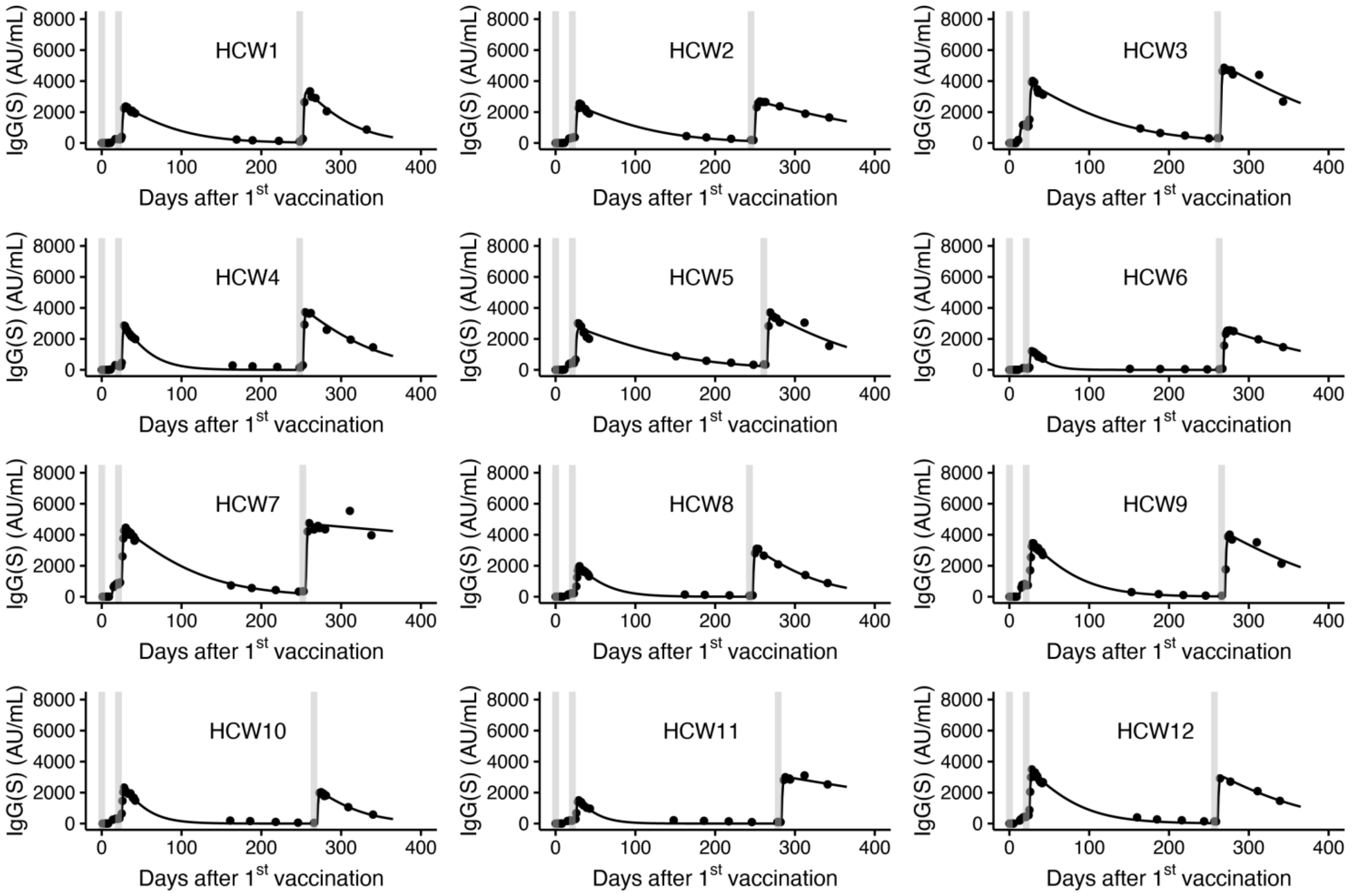


**Supplementary Figure 14. Calibrating primary and booster vaccine-elicited antibody dynamics:** Observed and best-fitted antibody titers are described for the 12 health care workers (HCWs) whose serum was sequentially sampled. The closed dots and solid curves indicate observed and reconstructed IgG(S) titers, respectively. The gray vertical lines correspond to the date of the primary two and booster vaccinations.

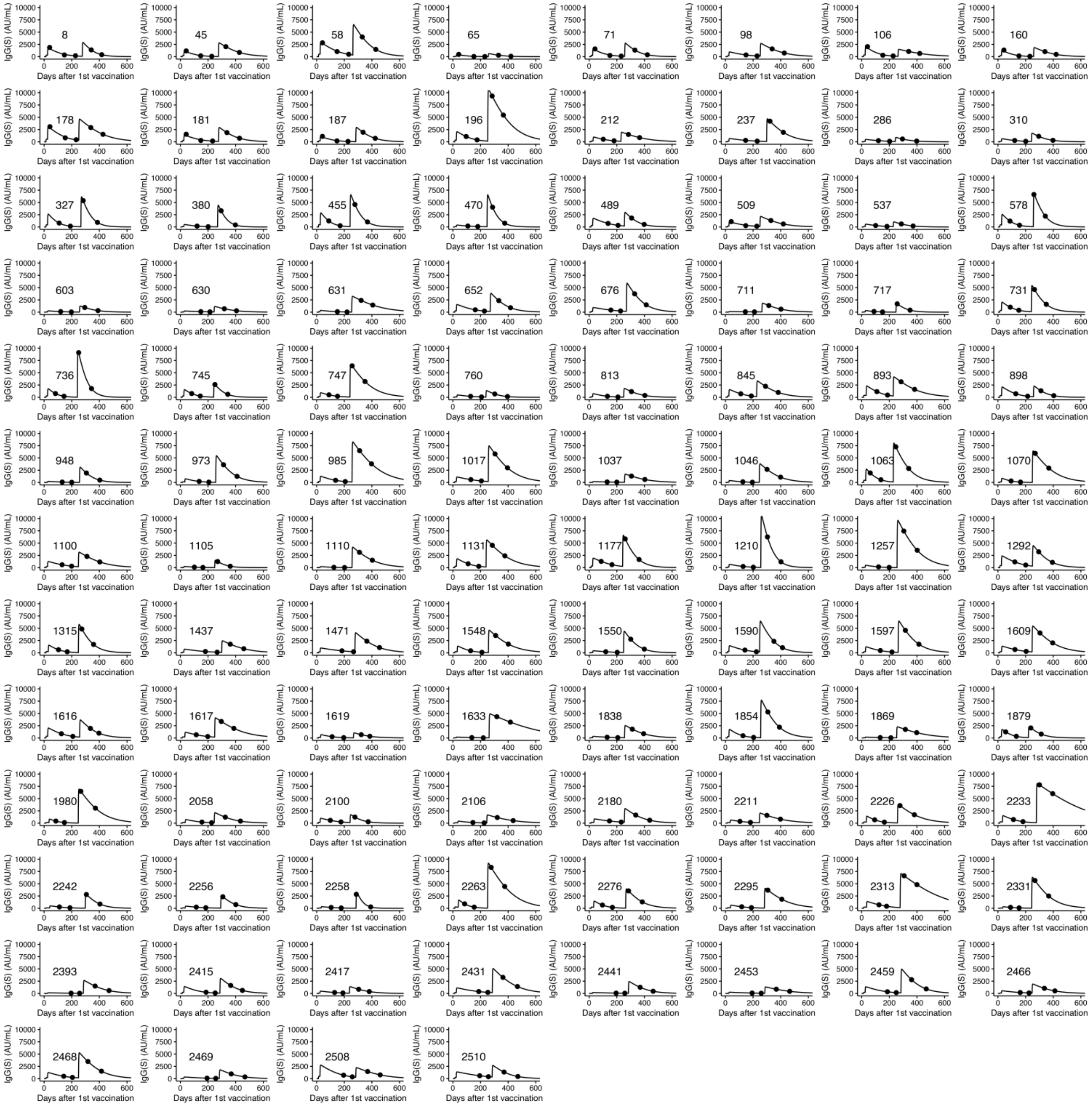


**Supplementary Figure 15. Reconstructed antibody titer trajectory for individual participants in the Fukushima vaccination cohort:** The estimated antibody titer for each individual participant (solid lines) along with the observed data (closed dots) are depicted using the best-fit parameter estimates. The curve of 100 was randomly selected for visualization because of the large number.

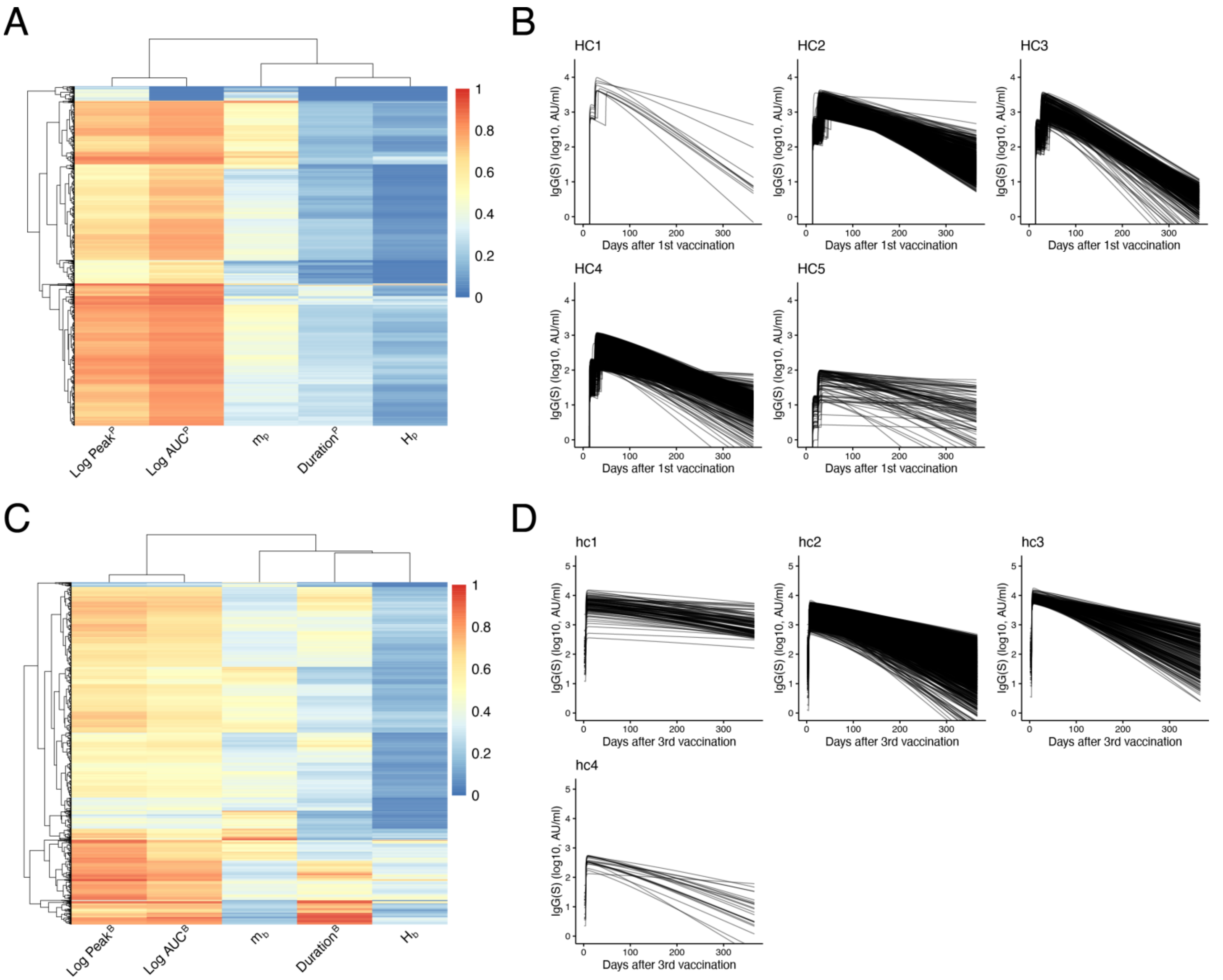


**Supplementary Figure 16. Unsupervised stratification of vaccine-elicited antibody response with hierarchical clustering: (A)** Heatmap of the five features (log10($\mathrm{Peak}^{\mathrm{P}}$), log10($\mathrm{AUC}^{\mathrm{P}}$), log10($m_p$), $\mathrm{Duration}^{\mathrm{P}}$, and $H_p$, normalized to 0-1), and hierarchical clustering are shown. **(B)** Reconstructed individual antibody dynamics after primary vaccination of 5 groups (HC1 to HC5) are represented. **(C)** Heatmap of the five features (log10($\mathrm{Peak}^{\mathrm{B}}$), log10($\mathrm{AUC}^{\mathrm{B}}$), log10($m_b$), $\mathrm{Duration}^{\mathrm{B}}$, and $H_b$, normalized to 0-1), and hierarchical clustering are shown. **(B)** Reconstructed individual antibody dynamics after booster vaccination of 4 groups (hc1 to hc4) are represented.

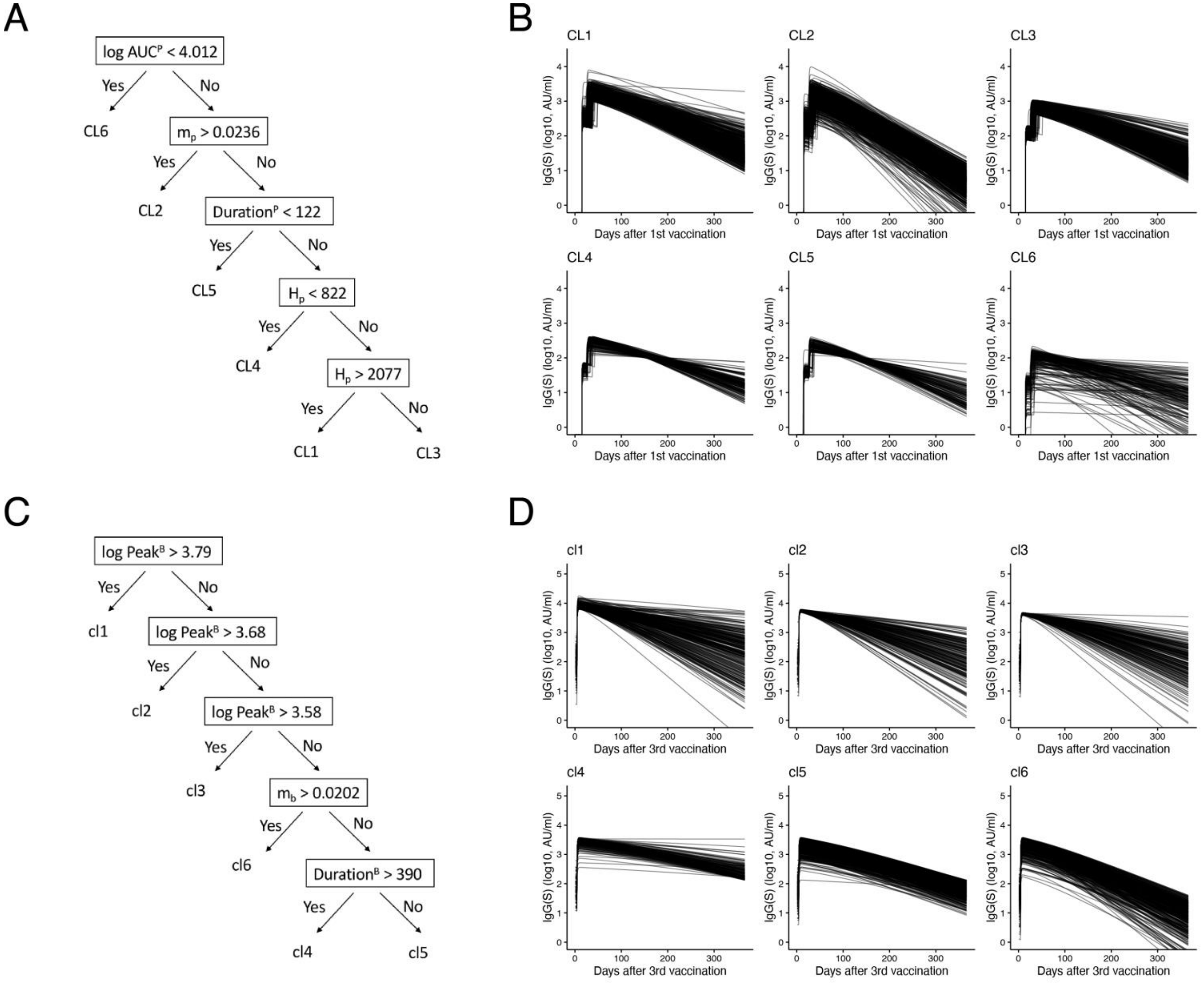


**Supplementary Figure 17. Unsupervised stratification of vaccine-elicited antibody response with a decision tree: (A)** Decision tree constructed with CLTree (https://github.com/dimitrs/CLTree) for five features (log10( $\mathrm{Peak}^{\mathrm{P}}$ ), log10( $\mathrm{AUC}^{\mathrm{P}}$ ), log10( $m_p$ ), $\mathrm{Duration}^{\mathrm{P}}$, and $H_p$ ) is shown. **(B)** Reconstructed individual antibody dynamics after primary vaccination of 6 groups (CL1 to CL6) are represented. **(C)** Decision tree constructed with CLTree (https://github.com/dimitrs/CLTree) for five features (log10($\mathrm{Peak}^{\mathrm{B}}$), log10($\mathrm{AUC}^{\mathrm{B}}$), log10($m_b$), $\mathrm{Duration}^{\mathrm{B}}$, and $H_b$,) is shown. **(B)** Reconstructed individual antibody dynamics after booster vaccination of 6 groups (cl1 to cl6) are represented.

### Supplementary Table 1. Estimated fixed and individual parameters for 12 health care workers

| Parameter or variable | Decay rate of antibody-secreting cells | Maximum *de novo* production of antibody by 1st vaccination | Delay of induction of antibody-secreting cells after 1st vaccination | Steepness at which induction increases with amount of mRNA after primary 2 dose | Amount of mRNA satisfying $P_i/2$ | Maximum *de novo* production of antibody by 2nd vaccination | Delay of induction of antibody-secreting cells after 2nd vaccination | Maximum *de novo* production of antibody by 3rd vaccination | Delay of induction of antibody-secreting cells after 3rd vaccination | Steepness at which induction increases with amount of mRNA after booster dose |
|---|---|---|---|---|---|---|---|---|---|---|
| **Symbol** | $\mu$ | $H_1$ | $\eta_1$ | $m_1$ | $K$ | $H_2$ | $\eta_2$ | $H_3$ | $\eta_3$ | $m_3$ |
| **Unit** | $\text{day}^{-1}$ | AU/mL | day | --- | $\mu g/0.5\text{mL}$ | AU/mL | day | AU/mL | day | --- |
| **Individual estimated parameters for $S1$ to $S12$** | | | | | | | | | | |
| $S1$ | 0.875 | 714.3 | 12.7 | 0.0304 | 28100 | 4833.5 | 3.9 | 7364.6 | 4.4 | 0.0361 |
| $S2$ | 0.875 | 795 | 13 | 0.0231 | 28100 | 4724 | 4.1 | 5075.8 | 4.5 | 0.013 |
| $S3$ | 0.875 | 2465.8 | 12.5 | 0.0207 | 28100 | 7370.2 | 3.8 | 9430.3 | 3.5 | 0.0151 |
| $S4$ | 0.875 | 1082.8 | 12.7 | 0.0602 | 28100 | 7063.7 | 3.9 | 7468.7 | 3.9 | 0.0244 |
| $S5$ | 0.875 | 935.2 | 12.7 | 0.0187 | 28100 | 5252.9 | 3.9 | 7001.8 | 4.3 | 0.0194 |
| $S6$ | 0.875 | 692.1 | 13 | 0.0869 | 28100 | 3807.3 | 4.1 | 5016 | 4.9 | 0.0173 |
| $S7$ | 0.875 | 1876.8 | 13.3 | 0.0234 | 28100 | 8646.4 | 4.2 | 8327.1 | 3.6 | 0.0027 |
| $S8$ | 0.875 | 729.3 | 13.9 | 0.0524 | 28100 | 4577.44 | 4.7 | 6318 | 4.2 | 0.0278 |
| $S9$ | 0.875 | 1992.9 | 13.6 | 0.037 | 28100 | 7421.13 | 4.5 | 7787.1 | 4.4 | 0.0177 |
| $S10$ | 0.875 | 1083.3 | 12.6 | 0.0578 | 28100 | 5512.7 | 3.8 | 4462.5 | 4 | 0.0386 |
| $S11$ | 0.875 | 806.4 | 13.5 | 0.0696 | 28100 | 4044.4 | 4.4 | 5519.3 | 4.1 | 0.0078 |
| $S12$ | 0.875 | 1119.2 | 12.6 | 0.0320 | 28100 | 6835.6 | 3.9 | 6027.6 | 4 | 0.0207 |
| **Population estimated parameters** | | | | | | | | | | |
| --- | 0.875 | 1097.7 | 13.0 | 0.0423 | 28100 | 5786.5 | 4.1 | 6472.4 | 4.1 | 0.0163 |
